# Conserved transcriptional reprogramming in nematode infected root cells

**DOI:** 10.1101/2025.09.05.674408

**Authors:** Maite Saura-Sanchez, Almudena Gómez Rojas, Melissa Deveux, Max Minne, Carolin Grones, Thomas Eekhout, Patricia Abril-Urias, Michiel Van Bel, Ruben Tenorio Berrio, Klaas Vandepoele, Carolina Escobar Lucas, Tom Beeckman, Bert De Rybel, Tina Kyndt

**Affiliations:** Ghent University, Department of Plant Biotechnology and Bioinformatics, Ghent, Belgium; VIB Center for Plant Systems Biology, Ghent, Belgium; Ghent University, Faculty of Bioscience Engineering, Department Biotechnology, Ghent, Belgium; Facultad de Ciencias Ambientales y Bioquímica, Universidad de Castilla-La Mancha, Toledo, Spain; VIB Single Cell Core Facility

**Author notes:** Laboratory of Cell and Developmental Biology, Cluster of Plant Developmental Biology, Department of Plant Sciences, Wageningen University, Droevendaalsesteeg 1, 6708 PB Wageningen, the Netherlands. equal contribution (Almudena Gómez Rojas, Melissa Deveux). Correspondence: Tina Kyndt, Bert De Rybel, Tom Beeckman, Carolina Escobar and Maite Saura-Sanchez.

## Abstract

Plant-parasitic nematodes are responsible for important annual losses in crop productivity worldwide^1,2^. Although the formation of feeding organs within the roots is essential for successful sedentary parasitism^3^, the molecular mechanisms underlying their development are poorly understood. This is partly because these organs originate from a limited number of root cells^4–7^, making difficult to capture the transcriptional reprogramming that occurs during the early stages of the infection. Here, we first developed a comparative host-pathogen framework to study the nematode infection process in *Arabidopsis* and rice. Using a cross-species single-cell transcriptomics approach, we identified a unique molecular signature in infected root cells and show that the cellular reprogramming during these early stages is highly conserved across both host-pathogen interactions. This transcriptional cell reprogramming is associated with stemness acquisition related to *de novo* organogenesis process. By cell-type specific gene regulatory network analysis, we identified AtATHB2/OsHOX28 as an evolutionary conserved and key regulator of the nematode infection process. Loss-of-function of this regulator in both species results in nematode resistance without affecting root growth. This discovery opens up new avenues for the development of sustainable nematode control strategies that could be translated across crop species.

## Introduction

Plant-parasitic nematodes are a threat to global agriculture^1^, causing an estimated 12% of all annual crop yield losses worldwide^2^. Among plant-parasitic nematodes, the endoparasitic root-knot nematodes (RKN; *Meloidogyne* spp.) are the most damaging group due to their wide host range and global distribution^8^. RKNs employ stylet-based feeding^3^ to withdraw nutrients and water from the host root cells^9^, hijacking the plant cellular machinery to develop highly specialized feeding structures known as giant cells^4,6,7,10^. These giant cells are accommodated inside a pseudo-organ called the gall, which is caused by the abnormal growth of the surrounding root tissue^5,11^. The formation of giant cells is indispensable for the nematodes to complete their life cycle, and detrimental to the development, fitness and reproduction capacity of the host, thereby reducing plant productivity^12–14^. As such, understanding how the formation of this organ occurs is crucial to develop efficient strategies to protect crop species against RNKs.

The ontogeny of galls and giant cells requires the recruitment of developmental pathways typically associated with transient pluripotent cell identities during root organogenesis or regeneration^15,16^. The spatial dynamics of gall and giant cell formation are intricately linked to the trajectory of the nematode within the root. Juvenile (J2) RKNs migrate intercellularly through cortical cells toward the vascular cylinder, where they settle permanently^17,18^. RKNs secrete effectors into a small number of selected cells^19^; inducing significant changes in their gene expression^20–22^ causing a reprogramming into giant cells. Ultimately, those cells show aberrant cell cycle activity^23^, and undergo drastic cellular changes. This process is accompanied by hypertrophy and hyperplasia of the cortical tissue^24^, forming visible galls. The intricate convergence of the nematode effectors, with plant cell-type-specific responses and dynamically regulated hormonal gradients^25^ orchestrates the structural and cellular modifications required for giant cell and gall formation.

Given the spatiotemporally complex nature of gall and giant cell development, bulk-RNA sequencing approaches failed to resolve the cellular heterogeneity responses at the molecular level. More targeted studies on laser micro-dissected giant cells have focused on developmental stages after the giant cells become morphologically distinguishable^26–30^, thereby missing the critical early molecular events driving their formation. Furthermore, a great number of studies have been focusing on *Arabidopsis thaliana*^31^, a non-natural host model species^32^. However, the putative transferability of this knowledge to applications in crop species is not always immediately obvious. Here, we addressed this set of limitations by applying a comparative cross-species scRNA-sequencing approach to resolve the spatiotemporal dynamics of gall and giant cell development in *Arabidopsis thaliana*, a well-characterised dicot model species and *Oryza sativa* (rice), a representative monocot and natural host crop.

### Nematode responsive cells show a unique spatiotemporal transcriptional signature

In order to pinpoint conserved molecular regulators during the early stages of RKNs compatible interaction, we characterized two diverged host-pathogen interactions using *Arabidopsis thaliana* (*Arabidopsis*) and *Oryza sativa* (rice) as host plants infected with *Meloidogyne javanica* and *Meloidogyne graminicola*, respectively. We conducted a time-course analysis of infection in both interaction systems, using acid fuchsin staining to visualize the nematode position within the roots (**Fig. 1a,b**). In *Arabidopsis*, we also monitored the expression of two known plant marker genes, *AtGATA23*^33^ and *AtPUCHI*^34^, early induced after nematode infection which were consistently expressed across all the evaluated time points (**Extended Data Fig. 1a,b**). Given the asynchronous nature of the infections, different stages of the infection were detected at every time point (**Fig. 1a,b**). We determined that 36 hpi for *Arabidopsis* and 42 hpi in rice, were the optimal timepoints for our analyses as we observed both a proportion of nematodes migrating within the root as well as a proportion already in the vascular cylinder (**Fig. 1a-d**), without presenting major morphological re-arrangements surrounding the nematodes as observed after 48 hpi (**Extended Data Fig. 1c,d).** At the selected timepoints, we also observed cells in the cortex undergoing cell division and/or expansion (**Fig. 1e,f**); morphological changes related to the accommodation of giant cells within the gall^35^. In summary, we have identified comparable developmental stages in both host-pathogen systems, representing a snapshot of the early stages of RKN infection.

**Fig. 1.**
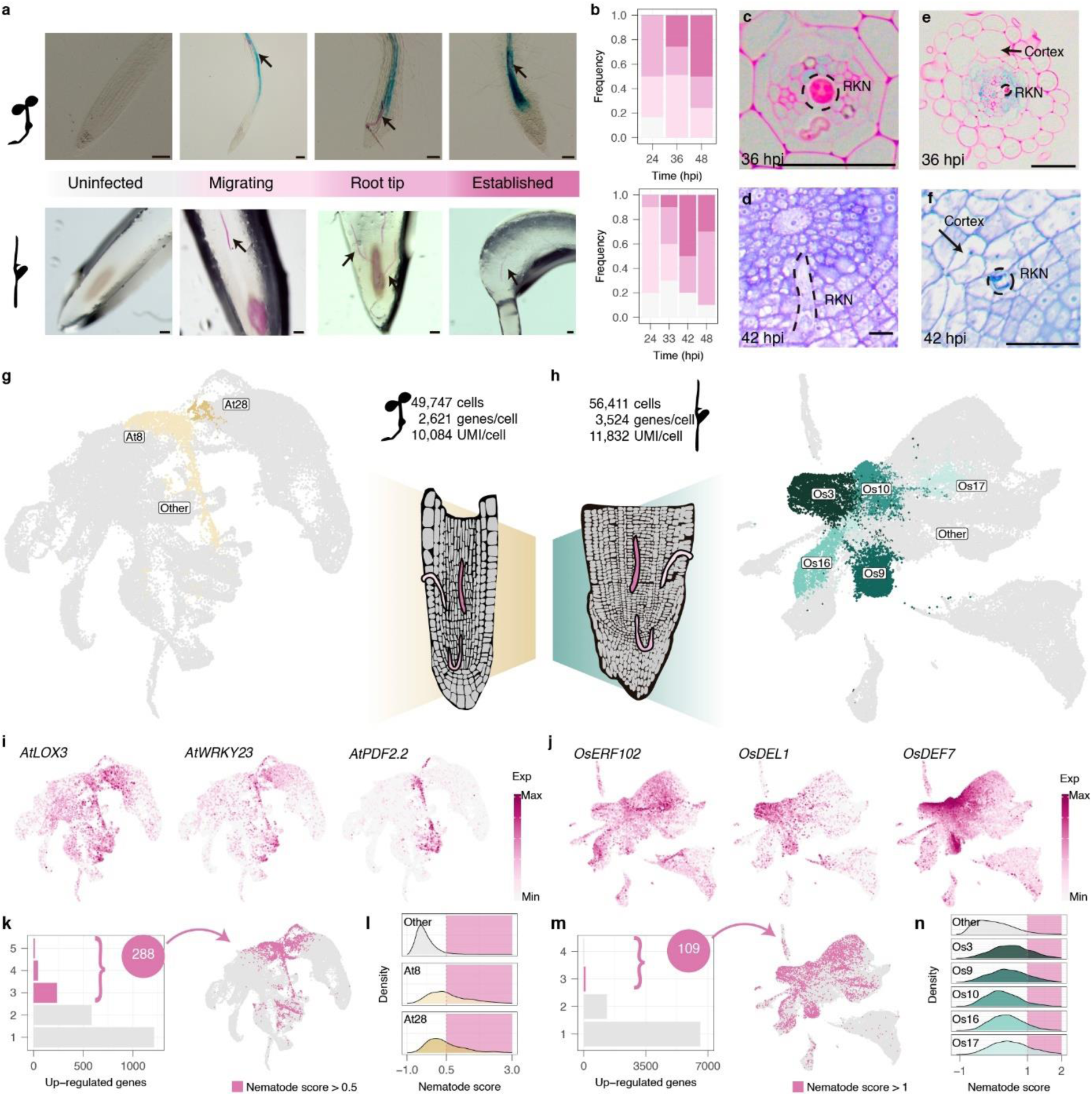
Nematode responsive cells group in unique “nematode clusters”. **a,b,** Evaluation of infection stages at different times during the early interaction of *Meloidogyne javanica* and *Arabidopsis thaliana* and *Meloidogyne graminicola* and *Oryza sativa*. **a,** Fuchsin stained roots representative of the different infection stages. Above, *Arabidopsis* roots; below, rice roots. **b,** Number of roots in every infection stage at the hours post infection (hpi) indicated. Scale bars, 100 μm **c,d,** Representative cross-section showing the presence of RKN head established in *Arabidopsis* roots at 36 hpi (**c**) and in rice roots at 42 hpi (**d**). Dashed lines mark RKN head. Scale bars, 100 μm. **e,f,** Representative cross-section showing morphological changes of cortex cells in *Arabidopsis* roots at 36 hpi (**e**) and in rice roots at 42 hpi (**f**). Dashed lines mark RKNs and arrows point towards modified cortex cells. Scale bars, 100 μm. **g,h,** Uniform manifold approximation and projection (UMAP) of merged mock and infected roots of *Arabidopsis* (**g**) and rice (**h**) datasets. Clusters enriched in cells from the infected samples are labelled and highlighted by non-grey colors. **i,j,** UMAPs showing the normalized expression levels of known RKN infection marker genes in (**i**) *Arabidopsis* and (**j**) rice datasets. **k-n,** Evaluation of nematode core genes expression in the scRNA-seq data. **k,m,** UMAP highlighting cells that surpass the threshold nematode score, given by the expression of genes constitutively up-regulated under nematode infection in previous studies in *Arabidopsis* (**k**) and rice (**m**) datasets. **l,n,** Distribution of values of nematode scores in the cells within the “nematode clusters” and the rest of the clusters (Other) of *Arabidopsis* (**l**) and rice (**n**) datasets.

We next profiled the spatiotemporal transcriptional changes during the very early steps of gall formation by conducting single cell RNA sequencing (scRNA-seq) on at least three independent replicates for both mock and inoculated root tips at 36 hpi in *Arabidopsis* and 42 hpi for rice (**Extended Data** Fig. 1e **and Supplementary Table 1**). Following raw data pre-processing and filtering (**Supplementary Table 1**), we retained 49.747 high-quality cells for the *Arabidopsis* dataset with an average of 2.621 genes per cell (**Fig. 1g and Supplementary Table 1**). For the rice dataset, 56.411 high-quality cells were retained for further analysis with an average of 3.524 genes per cell (**Fig. 1h and Supplementary Table 1**). A principal component analysis (PCA) (**Extended Data Fig. 1f,g**) and the Pearson correlation between samples (**Extended Data Fig. 1h,i**) showed that the main source of variability was not due to the biological replicates which were performed on different days and therefore, no additional batch correction was applied to the data. We next hypothesized that the cells undergoing transcriptional reprogramming contributing to giant cell and gall formation should be exclusively present in RKN infected samples and would likely form unique clusters. We thus selected the top clusters enriched with cells from the infected samples (clusters At8 and At28 for *Arabidopsis* and clusters Os3, Os9, Os10, Os16 and Os17 for rice) as potential clusters of interest (**Fig 1g,h and Extended Data Fig. 2a-d**). We assessed whether the cluster selection was biased by the difference in captured cells in mock versus infected samples by repeating the analysis with down-sampled datasets with a bootstrapping approach (**Extended Data Fig. 2e,f**). The cluster enrichment was consistent, justifying the use of the complete datasets for further downstream analysis.

Next, we verified the expression of marker genes in our single cell dataset which have earlier been reported to be induced during the early stages of sedentary nematode parasitism. Notably, in *Arabidopsis*, we observed *AtTDR1*^15^, *AtLOX3*^36^ and *AtPDF2.2*^37^ as markers of cluster At8; *AtARR5*^38^, *AtCEL1*^39^ and *AtERF109*^40^ as markers of cluster At28, and *AtWRKY23*^41^ as markers for both clusters (**Fig. 1i and Supplementary Table 1**). The orthologous genes of *AtERF109*, *AtDEL1*^42^ and *AtPDF2.2* in rice (Os09g0457900/ *OsERF102*, Os09g0457900/ *OsDEL1* and Os02g0629800/ *OsDEF7*, respectively) were also found as markers among the clusters Os3, Os9, Os10, Os16 and Os17 in the rice dataset (**Fig. 1j and Supplementary Table 1**). Additionally, we took advantage of available bulk whole genome transcriptomic datasets of nematode infection at different stages of RKN infection in rice and *Arabidopsis* (**Supplementary Table 1**). For each species, we identified a “nematode core set” of genes upregulated in infected versus uninfected roots in at least three studies (**Supplementary Table 1 and Methods**). Then we assigned a “nematode score” to every cell depending on the expression of this set of genes. For both species, cells with the highest “nematode score” were enriched in the selected clusters, as well in some of the neighbouring clusters (**Fig. 1l,n**). In summary, our findings strongly suggest that reprogrammed cells during RKN infection are present within the selected clusters of interest, hereafter referred to as “nematode clusters”.

### Transcriptional responses to nematode infection are conserved across species

In order to explore the similarities between both host-pathogen interactions during the early stages of gall and giant cell development, both datasets were integrated based on the expression of one-to-one orthologous genes (**Supplementary Table 2**). Using the same criteria as before, we looked for the clusters enriched in cells from the infected samples within the integrated dataset, and we identified clusters Int6 and cluster Int7 as the “nematode clusters” (**Fig. 2a and Extended Data Fig. 3a,b**). The cluster Int6 was mainly composed of cells originally belonging to “nematode clusters” from both species, including the cells from At8 and At28 in *Arabidopsis* and from Os10, Os16 and Os17 in rice (**Fig. 2b,c and Extended Data Fig. 3c**). The cluster Int7 contained most cells from the “nematode clusters” Os3 and Os9 of rice, as well as from the rice cluster Os21 and clusters At15 and At27 of *Arabidopsis* (**Fig. 2b,c and Extended Data Fig. 3c**). To further explore the transcriptional similarity between species, we computed pairwise rank correlations of the gene expression specificity index for one-to-one orthologous genes within the clusters^43^ (**Methods**). Notably, the correlation values for these clusters were higher than any other pairwise cluster comparisons, supporting the correspondence between the “nematode clusters” of both species (**Fig. 2d**).

**Fig. 2.**
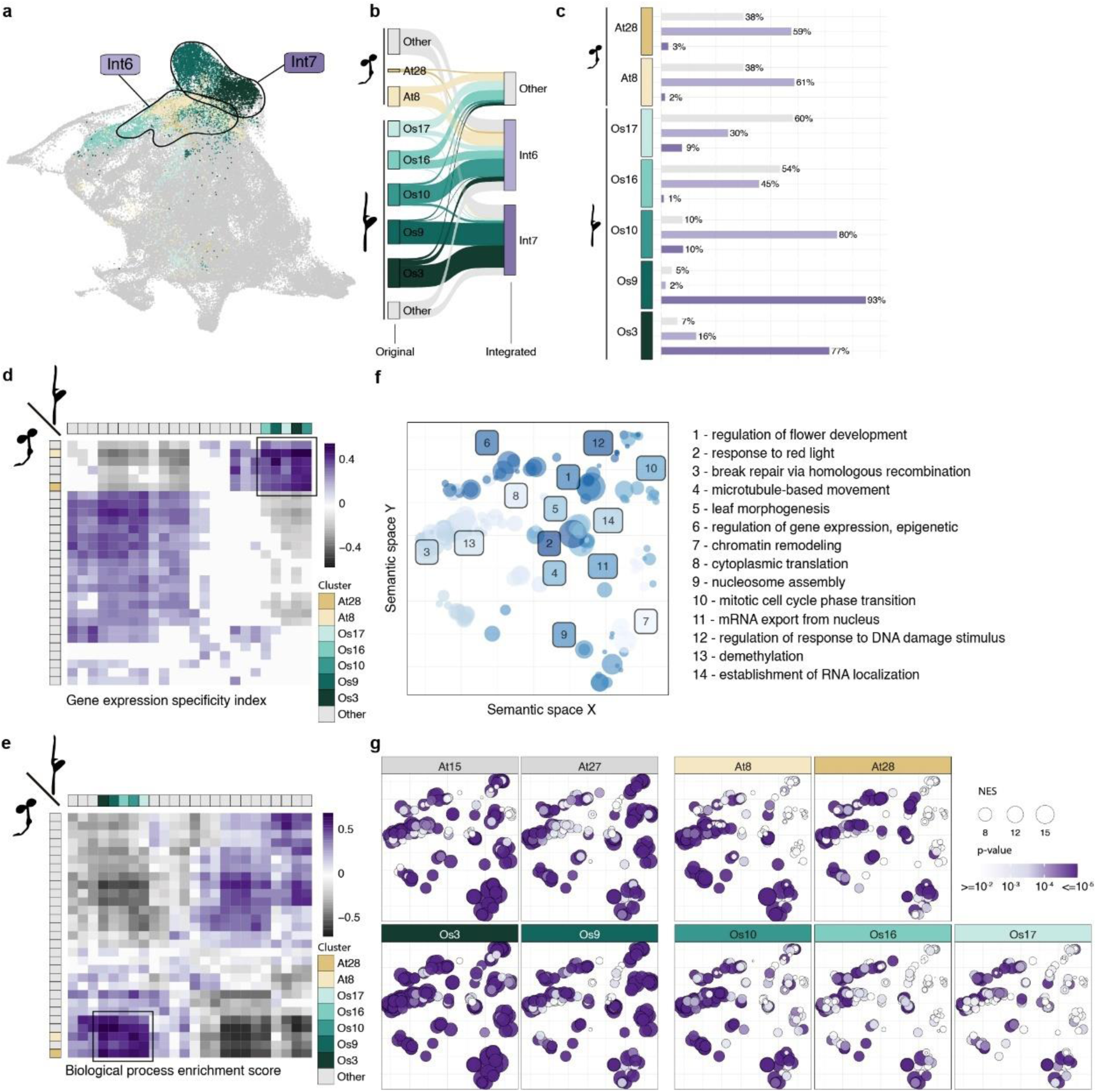
The “nematode clusters” are transcriptionally and functionally highly conserved between *Arabidopsis* and rice. **a,** UMAP of *Arabidopsis* and rice integrated dataset. Cells are colored by their original color on the UMAP analysis in Fig. 1. Circles surround cells of the nematode clusters Int6 and Int7 identified in the integrated dataset that are enriched in cells from the infected samples. **b,** Sankey diagram showing the correspondence between the cells in the original datasets and the integrated dataset. Only the cells that belong to “nematode clusters” either in the original or the integrated dataset are represented. **c,** Percentage of cells from the “nematode clusters” in the original datasets within the “nematode clusters” in the integrated dataset. **d,** Pairwise Spearman’s rank correlation of average gene expression between the clusters of *Arabidopsis* and rice datasets. Non-significant correlations are shown as coefficient zero. **e,** Pairwise Spearman’s rank correlation of normalized enrichment score (NES) of biological process (BP) GO terms, obtained by a simple sample gene set enrichment analysis (ssGSEA), between the clusters of *Arabidopsis* and rice datasets. Non-significant correlations are shown as coefficient zero. **f,** Semantic GO relationship of the top BP GO terms with the highest NES in the “nematode clusters”. **g,** Distribution of NES and p-values of the top BP GO across the “nematode clusters”, within the semantic space shown in **f**.

Comparisons of scRNA-seq datasets often only rely on one-to-one orthologous genes^44,45^. However, this approach is limited when analysing evolutionary distant species as, due to frequently duplication mechanisms during plant diversification, the one-to-one ortholog inference is often compromised obstructing the comparative analysis^46^. Furthermore, we observed a potential bias in such comparisons, as certain clusters showed an enrichment of differential expressed genes within the one-to-one ortholog subset (**Extended Data Fig. 4a**) which may exaggerate their similarity while overlooking features that represent other clusters. To overcome this limitation, we extended our analysis by transforming the aggregated gene expression per cluster into gene ontology (GO) biological processes (BP) terms with a single sample gene set enrichment analysis (ssGSEA)^47^. This approach considers all genes in the dataset of each plant species and ensures comparability across species^48^. Spearman correlation of the resulting normalized enrichment scores indicated high functional correlation of the “nematode clusters” between species (**Fig. 2e**). The semantic analysis of the top BP GO terms that define the “nematode clusters” were associated with key processes related to epigenetic regulation, RNA transport, cell division, and microtubule-based movement, which align with the extensive cellular reprogramming required during gall and giant cell formation (**Fig. 2f,g**). We also spotted distinct patterns of similarity within the “nematode clusters” (**Fig. 2g and Extended Data Fig. 4b**). Specifically, clusters Os10, Os16, and Os17 in rice displayed greater similarity to clusters At8 and At28 in Arabidopsis, while clusters Os3 and Os9 in rice showed stronger similarity to At15 and At27, respectively (**Extended Data Fig. 4b**). This last group showed a larger enrichment of BP GO categories related to cell division (**Fig. 2f,g**). These distinctions were consistent with the similarities between the transcriptional profiles observed earlier in the integrated dataset (**Fig. 2b)**. Together, our analysis suggests that the cellular responses to RKN infection are predicted to be functionally and transcriptionally highly conserved between rice and *Arabidopsis*.

### Nematode infection reprogrammed cells show heterogeneity in cell type and state

The spatiotemporal resolution of our single cell data offers the opportunity to dissect the cell-type specific contribution during the very early events of giant cell formation, as well as changes occurring in the neighbouring tissues, both forming the gall. In order to achieve this, we first predicted the cell type identity by taking advantage of reference root maps. For the rice dataset, we generated a reference map by carefully annotating the clusters of the mock samples using known cell-type-specific markers (**Extended Data Fig. 5a,b and Supplementary Table 3**). Additionally, we validated our annotations by mRNA *in situ* hybridization and HCR-FISH analyses for clusters lacking clear expression of known cell type markers (**Extended Data Fig. 5c-i**). For the *Arabidopsis* dataset, we used a previously published and experimentally validated *Arabidopsis* root atlas as reference^49^. This annotation enabled us to predict the cell type identity for the complete datasets (**Fig. 3a-d**). In *Arabidopsis*, two cell populations were overrepresented in infected roots, which we annotated as columella-like and QC-like cells (**Fig. 3b,e,f**). This suggests that during infection, some cells ectopically acquire a columella-like and QC-like identity. The cells annotated as columella-like were grouped in the “nematode cluster” At28 (**Fig. 3e**). We confirmed that known columella stem cell markers GLV5^50^, was indeed ectopically expressed outside the root apical meristem (RAM) context during nematode infection (**Fig. 3g**). The cells annotated as QC-like appeared exclusively in *Arabidopsis* infected roots, but not in the mock, and grouped in the “nematode cluster” *At8* (**Fig. 3f**). Also here, using the QC marker WOX5^51^, we confirmed the ectopic appearance of QC-like cells during the nematode infection process (**Fig. 3h**). In rice, the cell populations annotated as initials were highly overrepresented in the infected roots (**Fig. 3d**). Furthermore, after having assigned the cell cycle state to every cell using previous information on specific gene markers that allow to distinguish between the S and G2/M phase (**Extended Data Fig. 6a,b and Supplementary Table 3**), we found that the “nematode clusters” Os3 and Os9 are the ones predicted to contain cells actively dividing (**Fig. 3i and Extended Data Fig. 6b**). Interestingly, while cells expressing markers of these clusters are found during formative cell division in the RAM of uninfected roots, those markers are additionally expressed in cells surrounding the nematode in infected roots (**Fig. 3j and Extended Data Fig. 6c,d**). In the *Arabidopsis* dataset we found that clusters At15 and At27 grouped the proliferating cells as shown by their expression of known markers of S and G2/M phase of the cell cycle (**Extended Data Fig. 6b,e-g**). Taken together, these results suggest that a subset of cells are being reprogrammed during nematode infection to re-acquire division competence and obtain a meristematic state that in *Arabidopsis* resembles the transcriptomic signature of QC and columella stem cells. These represent clear hallmarks of *de novo* organogenesis associated with nematode infection for gall and giant cell formation.

**Fig. 3.**
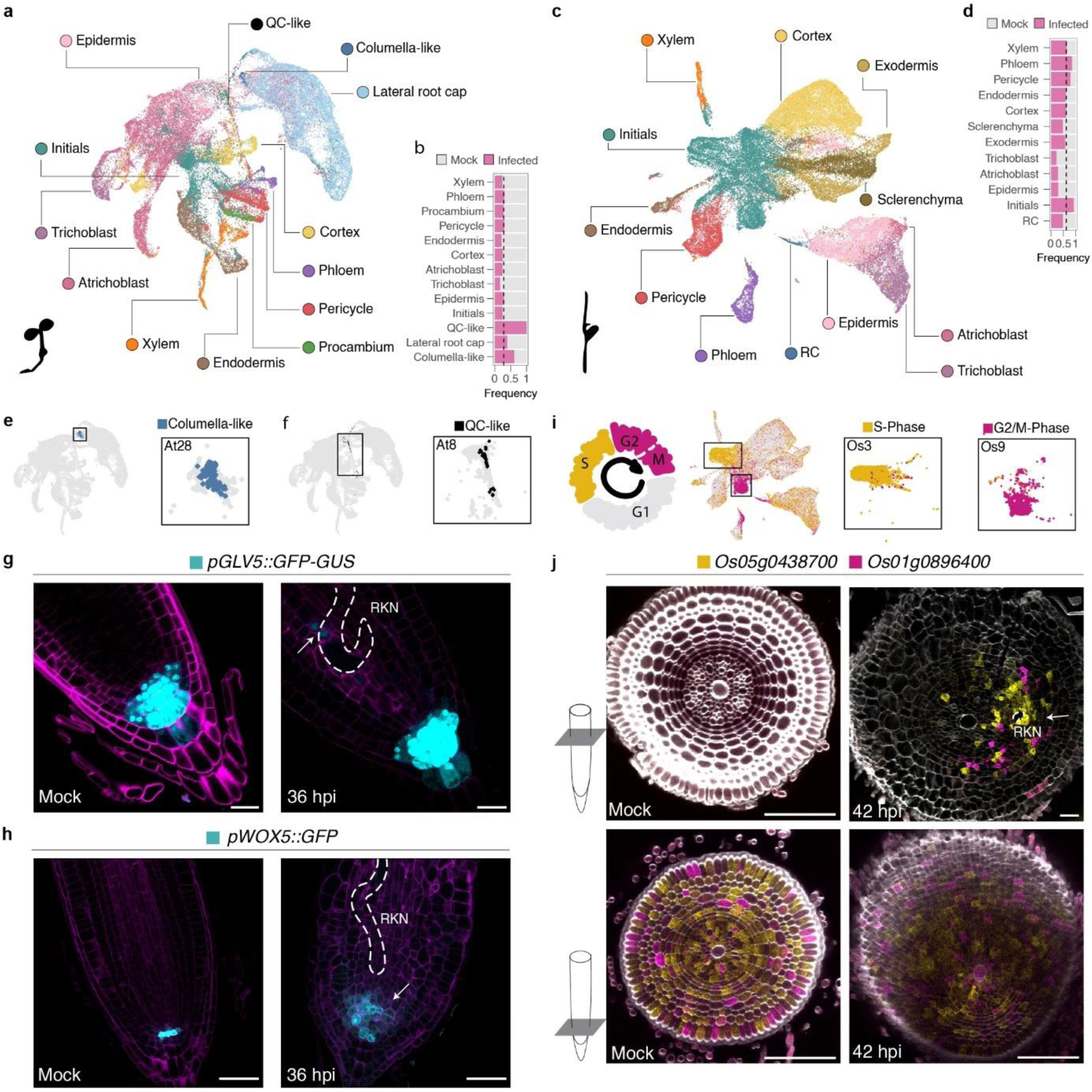
Reprogrammed cells within the “nematode clusters” are characterized by cell division competence and stemness transcriptional signatures. **a,** UMAP showing the predicted cell type identity in the *Arabidopsis* dataset. **b,** Frequency of cells from mock or infected samples by cell type in the *Arabidopsis* dataset. Dotted line indicates the expected frequency of infected cells. **c,** UMAP showing the predicted cell type identity in the rice dataset. **d,** Frequency of cells from mock or infected samples by cell type in the rice dataset. Dotted line indicates the expected frequency of infected cells. **e,** UMAP highlighting the cells predicted to be columella-like in the *Arabidopsis* dataset. **f,** UMAP highlighting the cells predicted to be QC-like in the *Arabidopsis* dataset. **g,h,** Confocal micrographs of *pGLV5::GFP* (**g**) and *pWOX5*::GFP (**h**) of *Arabidopsis* mock or infected roots at 36 hpi. Cell walls were stained with Direct Red 23 and represented in magenta. Scale bars, 20 μm (**g**) and 50 μm (**h**). LUT in GFP channel has been modified equally in mock and infected micrographs for better visualization. **i,** UMAP highlighting the cells predicted to be S-phase and G2/M-phase in the rice dataset. **j,** Fluorescence RNA *in situ* hybridization showing the transcript localization of marker genes from clusters Os3 and Os9. The confocal micrographs show cross sections of rice mock roots (left) or infected roots at 42 hpi (right). Cell walls were stained with calcofluor white and represented in grey scale. Scale bars, 100 μm. RKN: root-knot nematode. Arrows indicate fluorescence ectopic signal. Dashed white lines indicate nematodes.

The nematode clusters were predicted to contain a mix of cell type identities in both Arabidopsis (**Fig. 4a**) and rice (**Fig. 4b**) datasets. To link these cells with their localization during the nematode infection process, we selected specific and conserved markers (**Fig. 4c**) and explored their predicted patterns of expression during the early infection stages (**Fig. 4d**). Some marker genes such as *OsMYB30* and *OsDOF22* showed a broad induction in multiple cell types including cortex, endodermis, pericycle and vascular tissues in the vicinity of the nematode infection sites **(Fig. 4e,f)**, other markers such as *OsDEF7* showed more specific induction in cortical cells **(Fig. 4g).** Of even higher interest are evolutionary conserved markers of the infection process as these could act as general markers of the nematode infection process in a variety of plant species. One of the top candidate genes predicted to be specifically induced in the nematode clusters in both Arabidopsis and rice was *AtEXPANSIN4* (expressed clusters At8 and At28) and its ortholog in rice *OsEXPA7* (with higher expression in cluster Os17) (**Fig. 4h,i**). We validated their expression pattern during the infection process and found that both *AtEXPA4* and *OsEXPA7* were specifically expressed in cells surrounding the nematode in their respective host-pathogen interaction **(Fig. 4h,i)**. Notably, expression of *OsEXPA7* around the RKN was found in a region located between the phloem and xylem poles containing the inner cortex, endodermis and pericycle cells, where we observed that giant cells typically form in later developmental stages **(Fig. 4h,j)**. Also, the expression of *AtEXPA4* was observed at 4 dpi on binucleated cells surrounding the nematode established in the vascular cylinder (**Fig. 4k**). We therefore identified conserved orthologous genes specifically expressed in the precursors of the giant cells.

**Fig. 4.**
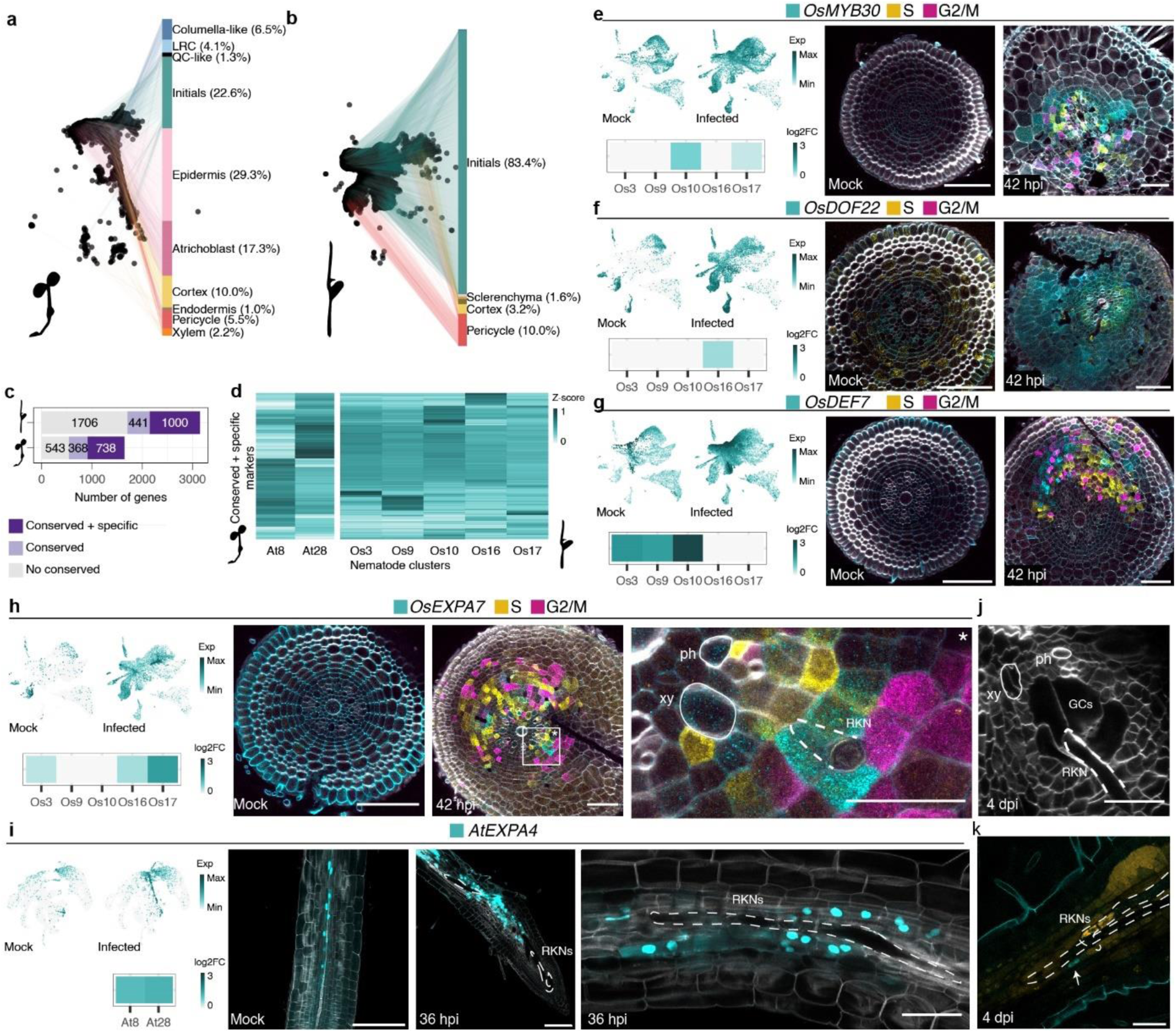
The “nematode clusters” contain the reprogrammed cells located in the tissues surrounding the nematode infection sites as well as the giant cell initials. **a,b,** Frequency of predicted cell types within the “nematode clusters” in *Arabidopsis* (**a**) and rice (**b**). Cell types with more than 1% of representation are labelled. **c,** Number of genes that are positive marker genes of the “nematode clusters” in *Arabidopsis* and rice. Genes that belong to orthogroups represented in the markers of both species are labelled as conserved. When additionally are highly specific to the “nematode clusters” are labelled as conserved+specific. **d,** Heatmap representing the gene expression profiles of the conserved+specific marker genes across the “nematode clusters”. Values represent the Z-score of gene expression considering all the clusters for every dataset. Left, data relative to *Arabidopsis*; right, data relative to rice. **e-i,** Gene expression profiles of the genes within the conserved+specific marker genes *OsMYB30* (**e**), *OsDOF22* (**f**), *OsDEF7* (**g**), *OsEXPA7* (**h**) and *AtEXPA4* (**i**). For every panel, UMAP showing the normalized expression of the gene and heatmap showing the log2FC of the marker genes across the “nematode clusters”. **e-h,** Fluorescence RNA *in situ* hybridization showing the transcript localization of the indicated genes. The confocal micrographs show cross sections of rice mock roots (left) or infected roots at 42 hpi (right). Asterisk marks the region zoom in. Cell walls were stained with calcofluor white and represented in grey scale. Scale bars, 100 μm. **i,** Confocal micrographs of *AtEXPA4::GFP-GUS* of *Arabidopsis* mock or infected roots at 36 hpi. Cell walls were stained with Direct Red 23 and represented in grey scale. Scale bars, 100 μm, 100 μm and 50 μm. **j,** Confocal micrograph shows a cross section of rice infected roots at 4 dpi containing formed giant cells. Cell walls were stained with calcofluor white and represented in grey scale. Scale bar, 50 μm. **k,** Confocal micrograph of *AtEXPA4::GFP-GUS* at 4 dpi. The arrow indicates binucleate cell. Nucleic acids were stained with SYTOX^TM^ ORANGE and represented in orange. Scale bar, 50 μm. RKN: root-knot nematode; xy: xylem; ph: phloem; GCs: giant cells. Dashed white lines indicate nematodes.

These findings confirm that our approach effectively captured the cells undergoing reprogramming; characterized by active division and the acquisition of a meristematic state. Both processes are thus likely required for gall and giant cell formation in later stages.

### Cell type-specific gene regulatory networks are rewired to reprogram cells towards giant cell and gall formation

To better understand the regulatory mechanisms underlying gall and giant cell formation, we analysed cell type-specific gene regulatory networks (GRNs) that could contribute to the reprogramming of cells identified in the “nematode clusters”. Using the gene regulatory network inference algorithm MINI-EX^52,53^ (**Methods**), we identified regulons (transcription factors (TFs) and their target genes) within each cell type that were specific to mock or infected conditions, as well as those shared between both treatments (**Fig. 5a-c and Supplementary Table 4**). In order to evaluate the uniqueness of the TFs for the mock and infected regulons (green and magenta bars in **Fig. 5b,c**), we next evaluated all regulons the TF is involved in across all cell types and plotted the proportion of mock and infected regulons (**Fig. 5d**). These results illustrate that most TFs operate in multiple regulons, not specific to the infection process.

**Fig. 5.**
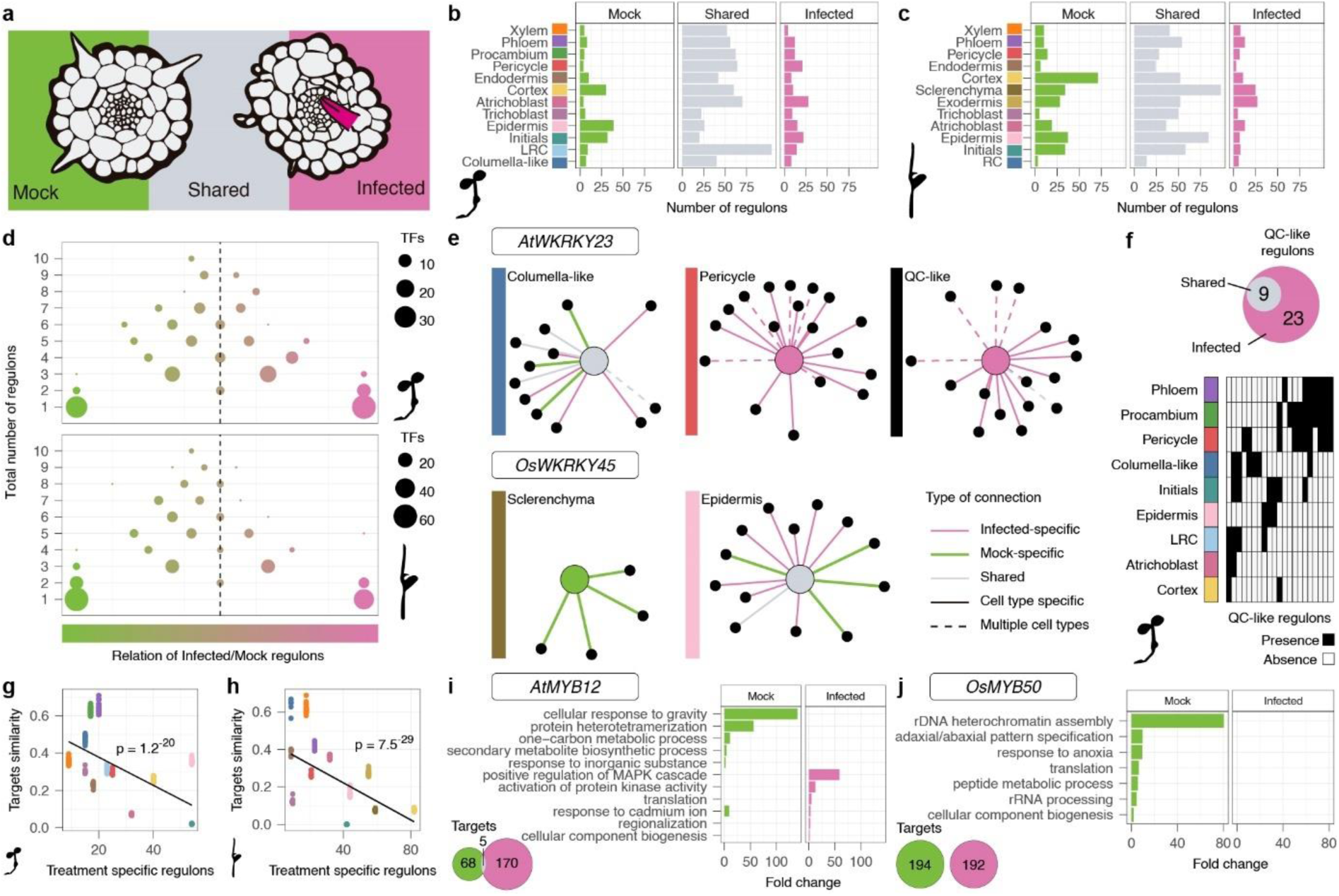
Cell-type-specific gene regulatory networks are rewired during nematode infection. **a-c,** Distribution of identified regulons across specific cell types in mock roots, infected roots or both treatments, referred as shared regulons, in *Arabidopsis* (**b**) or rice (**c**) datasets. **d,** Global behavior of transcription factors (TFs) that are mock specific (green bars in **b** and **c**) or infected specific (pink bars in **b** and **c**) in a given cell type, across the whole landscape of regulons. Dot size represents the number of TFs with the same behavior. Gradient color represents the treatment specificity of the TF taking in account all the regulons that controls across all the cell types in the data. **e,** Example of WRKY TFs that are identified as regulons. Above, *AtWRKY23* in the *Arabidopsis* dataset. Below, *OsWRKY45* in the rice dataset. Black dots represent target TFs that are also identified as regulon of the indicated cell type. **f,** Overlapping of TFs identified as regulons in the QC-like cell population and TFs identified as regulons in other cell types in *Arabidopsis*. **g,h,** Correlation between the Jaccard index of targets similarity of TFs that are shared in given cell type for both mock and infected treatments (grey bars in **b** and **c**) and the number of treatment specific regulons in the same cell type (green + pink bars in **b** and **c**). The p-value of the correlation is shown in the plot. **i,j,** Biological process enrichment of the targets of shared regulons governed by MYB TFs in initial cells either in mock or infected roots. **i,** *AtMYB12* in *Arabidopsis* (**i**) and *OsMYB50* in rice (**j**).

In other words, these results support the notion that nematode infection recruits or inhibits existing plant TF networks. Within regulons that are treatment-specific in at least one cell type, we found an enrichment of the same two orthogroups representing WRKY and ERF family members in both species (**Extended Data Fig. 7a-h and Supplementary Table 4**). Members of these TF families have been extensively linked to RKN infection ^10,54–60^. Our data predicts AtERF115 to be active in phloem tissue exclusively in infected roots (**Extended Data Fig. 7g**), in line with its putative role in gall organogenesis^61^. For the WRKY TF family this was exemplified by *AtWRKY23*, a shared regulon in the columella, that is ectopically activated in the pericycle and the QC-like population under RKN infection (**Fig. 5e and Extended Data Fig. 7g**), aligning with its known role in feeding site establishment^41^. In rice, *OsWRKY45*, a transcription factor associated with defense responses against RKNs^62^, was identified within this family and predicted to be repressed in sclerenchyma tissue under infection (**Fig. 5e and Extended Data Fig. 7h**). Another particularly striking example of ectopic expression was the emergence of cells with QC-like identity at infection sites in *Arabidopsis* (**Fig. 3f,h**). Since no QC cells were recovered in our mock root tip dataset, we compared the regulons of these ectopic QC-like cells with previously reported QC regulons^52^. Most of the regulons identified in these cells were distinctive to the nematode infection and largely overlapped with regulons identified in the pericycle and procambium tissues (**Fig. 5f**). This hints towards the involvement of these tissues in the formation of the QC-like cell population and stemness acquisition during RKN infection.

To further explore the global transcriptional rewiring landscape during nematode infection, we next analysed whether the TFs from the shared regulons within cell types (grey bars in **Fig.5b,c**) controlled similar downstream targets or whether this was rewired under nematode infection. We found that the overlap between the targets of TFs that are common to both mock and infected treatment decreased with the number of treatment-specific regulons within the cell type (**Fig. 5g,h, Extended Data Fig. 7i-l and Supplementary Table 4**). For instance, the TFs *AtMYB12* and *OsMYB50* were ranked highest among shared regulons in *Arabidopsis* and rice initial cells, respectively (**Extended Data Fig. 7m,n**). However, their target genes differed completely between mock and infected conditions, with each set enriched for distinct biological functions (**Fig. 5i,j**). These results suggest that the early stages of nematode infection involve massive transcriptional reprogramming by both changes in cell type-specific activation or repression of GRNs (**Fig. 5d-f**), and at the same time extensively rewiring of the existing ones (**Fig. 5g-j**).

### AtATHB2/OsHOX28 is required for root nematode infection

Given our approach successfully pinpointed cells undergoing transcriptional reprogramming during gall and giant cell development, along with novel marker genes and known transcription factors involved in RKN responses, we next determined whether novel conserved functional regulators required for gall formation could also be identified. We focused on those genes encoding TFs that were specifically expressed in the “nematode clusters” and ranked as cell type-specific regulons by MINI-EX (**Fig. 6a**). To ensure the selection of transcription factors was highly restricted to the nematode infection while minimizing potential effects on other developmental processes, we retained only those regulons that were active exclusively under infected conditions. In order to avoid technical artefacts, we discarded genes differentially expressed after protoplast isolation compared to dissected root tips. We generated this data by bulk RNA-seq for rice (**Supplementary Table 5 and Methods**) and used existing lists for Arabidopsis^63^. We finally only considered orthologous genes that met these criteria in both species. These stringent selection criteria (**Fig. 6a**) identified a small subset of three orthogroups representing ERF, MYB and homeodomain (HD)-ZIP transcription factors (**Supplementary Table 5**). The orthologous class II HD-ZIP pair *AtATHB2/OsHOX28* emerged as the most promising conserved candidate to specifically regulate the nematode infection process as it showed a clear one-to-one orthology between Arabidopsis and rice and was detected in the QC-like population in *Arabidopsis* (**Supplementary Table 5**). Both AtATHB2 and OsHOX28 have been linked to auxin signalling, a hormone crucial during gall formation^64–67^. In rice, OsHOX28 and OsHOX1 redundantly control local distribution of auxin during tiller angle establishment^68^, while in *Arabidopsis*, AtATHB2 maintains cell stemness by interfering with auxin signalling, together with its paralogs AtHAT3 and AtATHB4^69^. In our analysis, the AtATHB2 regulated network was ranked as an important regulon in the procambium and QC-like cells (**Fig. 6b**). Its paralog in rice, OsHOX28, was identified as a regulon in the cortex layer during nematode infection (**Fig. 6c**). Both were found as markers of the “nematode clusters” (Fig. 6d,e). We first validated these predictions by monitoring the translational reporter *pATHB2::ATHB2-GUS* in *Arabidopsis.* In uninfected roots, the GUS signal distributed uniformly within the vascular cylinder in the region close to the root meristem (**Fig. 6f**). In infected roots we observed an additional patchy signal in the elongation zone, probably coinciding with establishing nematodes (**Fig. 6f**). In rice, *OsHOX28* transcripts were weakly detected in vascular cells of uninfected roots, particularly in phloem companion cells (**Fig. 6g**). Upon infection, *OsHOX28* expression expanded ectopically in the infection zone to multiple tissues, including the cortex, pericycle and endodermis (**Fig. 6g**).

**Fig. 6.**
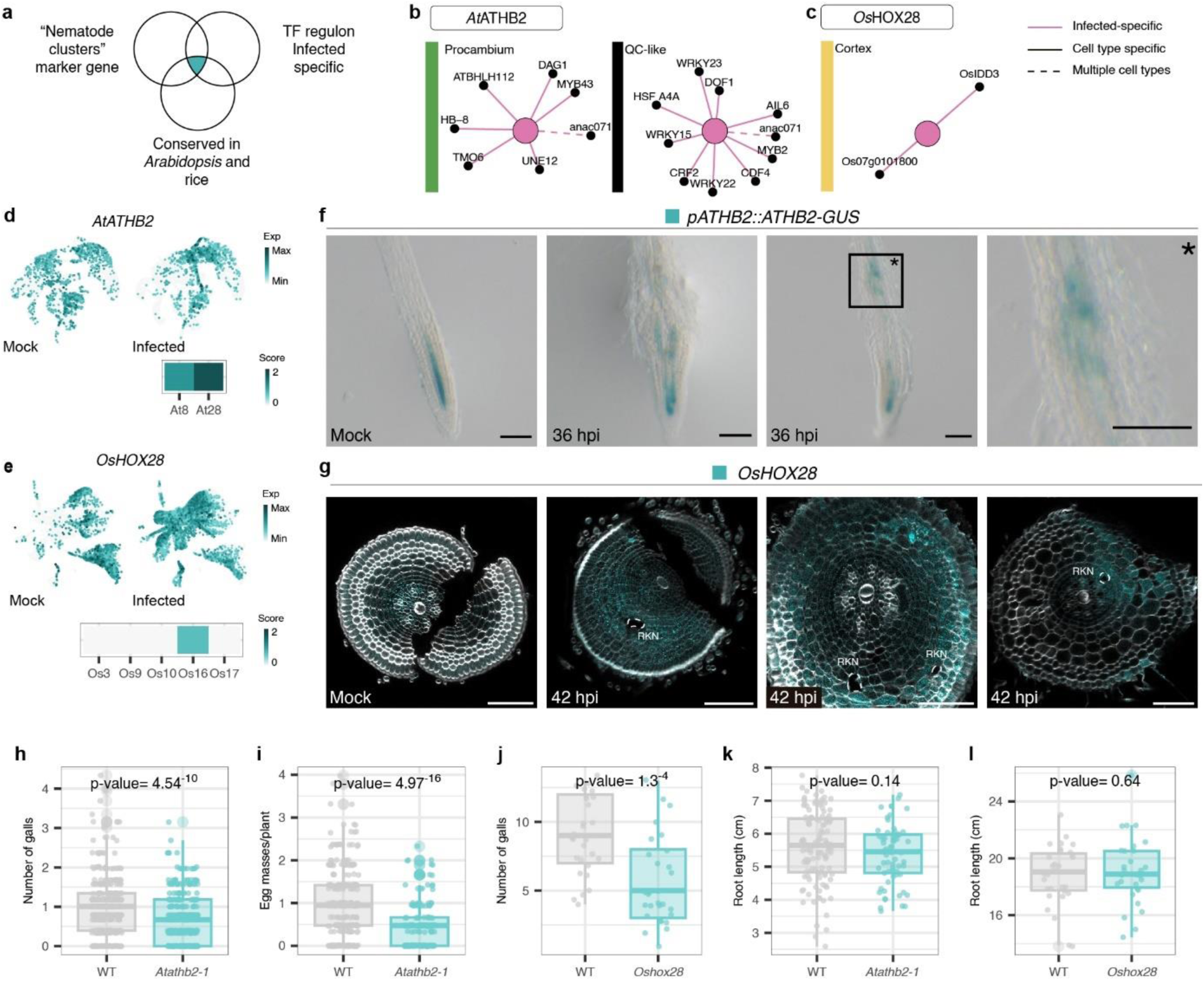
AtATHB/OsHOX28 is required for nematode gall formation. **a,** Criteria for gene selection **b,c,** Predicted networks for *At*ATHB2 (**b**) and *Os*HOX28 (**c**). Black dots represent target TFs that are also identified as regulon of the indicated cell type. **d,e,** UMAP showing the normalized expression of the gene and heatmap showing the score (pct.1/pct.2 * avgFC) of the gene across the “nematode clusters” for *AtATHB2* (**d**) and *OsHOX28* (**e**). **f,** Representative *Arabidopsis* roots showing the expression of *pATHB2::ATHB2-GUS* at 36 hpi during *Meloidogyne javanica* infection, as monitored by histochemical detection of GUS (blue substrate). Asterisk marks the region zoom in. Scale bars, 100 μm. **g,** Fluorescence RNA *in situ* hybridization showing the transcript localization of *OsHOX28*. The confocal micrographs show cross sections of rice mock roots or infected roots at 42 hpi. Cell walls were stained with calcofluor white and represented in grey scale. Dashed lines indicate the position of RKNs. The brightness of the whole images has been equally edited for better signal visualization. Scale bars, 100 μm. **h,** Number of galls per plant in *Arabidopsis* WT (n = 387) and *Atathb2-1* (n = 329) plants at 10 dpi. **i,** Number of egg masses per plant in *Arabidopsis* WT (n = 291) and *Atathb2-1* (n = 180) plants at 45 dpi. **j,** Number of galls per plant in rice WT (n = 28) and *Oshox28* (n = 29) plants at 7 dpi. **k,** Root length of *Arabidopsis* WT (n = 116) and *Atathb2-1* (n = 65) plants. **l,** Root length of rice WT (n = 30) and *oshox28* (n = 30) plants under control conditions. The boxplots show the median as a solid line, the hinges representing the 25th and 75th percentiles, and the whiskers that extend up to a maximum of 1.5 times the interquartile range from the hinges. Individual observations are represented by colored points. Exact p-values are shown for a two-sided t-test.

Having established that expression of both *AtATHB2* and *OsHOX28* orthologs is indeed induced during the nematode infection process in zones relevant to nematode infection and gall formation for the respective species, we next evaluated whether these regulators could potentially also play a functional role in the infection process. We thus infected an *Atathb2-1* mutant^70^ in *Arabidopsis* and an *Oshox28* mutant in rice with *M. javanica* and *M. graminicola*, respectively, and compared the number galls that developed on mutant roots compared to wild type roots. In both species, we observed an impaired of RKN infection **(Fig. 6h-j**) with more than 30% reduction in the number of nematode-induced galls per plant (**Fig. 6h,i)**. In *Arabidopsis*, we additionally analysed the reproduction capacity of the nematodes that was reduced up to a 50% in the *Atathb2-1* mutant compared to the wild type (**Fig. 6i**). Most important for the potential application of this conserved regulator, we did not detect developmental defects related to root growth due to the mutation (**Fig. 6k,l**). These results indicate that *AtATHB2*/*OsHOX28* is required for gall development in multiple plant-nematode interactions and can be used to engineer resistant plants to RKN infection.

## Discussion

In a broad range of host plants, parasitism by root knot nematodes relies on the formation of giant cells as feeding structures within the roots^10^. This intimate endoparasitic relationship challenges the development of effective pest control strategies in crop species, but also presents an opportunity to engineer loss of susceptibility^71^ by targeting key developmental regulators of gall and giant cell formation. Although this approach has been proposed for decades^72^, identifying early regulators of this organogenetic has been hampered by the dynamic and asynchronous nature of the nematode infection and the lack of early marker genes specific to this process. Here, we addressed these challenges by capturing a snapshot of the early transcriptional events leading to giant cell and gall formation using single cell RNA-sequencing in plant roots. Our results revealed massive transcriptional reprogramming at the tissue level and dynamic changes to the underlying gene regulatory network (**Fig. 5**). We were able to detect a distinct population of cells which we refer to as the “nematode clusters” which is reprogrammed during nematode infection and is characterized by changes in the cell cycle state and cell type identity (**Figs. 1, 3 and 4**). Among these, a subpopulation putatively corresponding to giant cell initials is characterized by the expression of *AtEXPA4/OsEXPA7* which we identified as a unique and evolutionary conserved marker of the process (**Fig. 4i-l**). As such, *AtEXPA4/OsEXPA7*, along with other co-expressed genes in our dataset, will serve as markers for the community to track and further understand the early events during RKN infection in multiple plant species. Despite the high resolution of our data, only very few cells express this unique marker in the infected roots. This confirms that bulk RNA-seq at this early stage of infection would fail to uncover the transcriptional changes associated with this organogenesis process.

The *de novo* organogenesis processes associated with giant cell formation have been extensively studied in *Arabidopsis* and dicot crops like tomato. RKNs recruit plant developmental pathways associated with post-embryonic organ formation to induce transient pluripotency for gall organogenesis^16^. Our dataset captured the transcriptional reprogramming associated with this organogenesis. Indeed, specific cell populations associated with the RKN infection process were found to be transcriptionally similar to QC stem cells (**Fig. 3**). Additionally, we observed that markers typically restricted to columella stem cells in uninfected roots were ectopically expressed during RKN infection (**Fig. 3**). This novel finding can be directly linked to the process of regeneration, where cells in the vascular tissue start expressing QC and columella marker genes during their re-specification soon after root damage^73^, a process known to be coupled with gall formation^40,61^.

In contrast to dicot species, the molecular mechanisms of gall organogenesis in monocots remains largely unknown. In this study, we have used the agriculturally relevant interaction between rice and *Meloidogyne graminicola* as a model system^74,75^. Our work provides novel insights into the early transcriptional events of this interaction, with extensive validation at unprecedented cellular resolution (**Figs. 3 and 4**). More importantly, we show that the early responses to RKN infection are highly similar between monocots and dicots (**Fig. 2**). This is unexpected given the phylogenetic distance between these species and the fact that cell type specific transcriptomes have been show to not be so equivalent^45^. Our results show that several executors of giant cell and gall formation are conserved between *Arabidopsis* and rice (**Fig. 4**), providing a foundation for translating findings related to RKN infection from dicots to monocots. Moreover, several genes we found to be involved in the early stages of RKN infection have been previously linked to the formation of syncytium feeding structures induced by plant-parasitic cyst nematodes (e.g. *AtEXPA4*^76^, *AtWKRY23*^41^*, AtPDF2.2*^37^*, AtERF109*^77^). Hence, our results could potentially be transferred to other plant-nematode interactions.

Identifying the developmental regulators of plants which get coopted during giant cell and gall organogenesis is instrumental to engineer a durable type of nematode resistance in crops which do not form feeding structures despite growing in RKN infested soils. By comparing the tissue-specific gene regulatory networks between RKN infected and uninfected roots, we identified known TFs related to pathogenesis and nematode responses (e.g. ERFs and WRKYs) in both species (**Fig. 5**). Moreover, our results suggest that the GRNs are extensively rewired in the sense that the same TFs regulate alternative target genes in the infected roots as compared to uninfected (**Fig. 5**). This observation is aligned with the notion that RKN susceptibility relies on hijacking existing developmental pathways and offers an explanation why targeting TFs to achieve RKN resistance is usually accompanied with pleiotropic effects on root architecture reducing plant fitness. The high spatiotemporal resolution and cross-species character of our approach, however, allowed us to narrow our selection to those TFs specific to the RKN infected state. We identified a conserved orthologous gene pair *AtATHB2/OsHOX28*, that encodes a class II HD-ZIP TF, with a crucial role in RKN parasitism in both *Arabidopsis* and rice. Indeed, *Meloidogyne* spp. infection and reproduction was severely impaired in *Atathb2-1* and *oshox28* loss of function *Arabidopsis* and rice mutant lines, respectively (**Fig. 6**). While other functions for AtATHB2 and OsHOX28 only resulted in phenotypes in association with other HD-Zip II TFs due to genetic redundancy^68–70^, here we show that the knock-out of AtATHB2 or OsHOX28 alone is sufficient to restrict RKN gall formation without affecting root growth (**Fig. 6**). As such, AtATHB2/OsHOX28 is a prime candidate to engineer RKN resistant plants in both monocot and dicot crop species.

## Acknowledgements

We would like to thank Monica Carabelli for sharing *pATHB2::ATHB2-GUS* and *athb2-1 Arabidopsis* seeds; Miguel Moreno-Risueño for sharing *pWOX5::erGFP* seeds; Lien de Smet for maintaining *M. graminicola* cultures and providing technical assistance; Marnik Vuylsteke, Jessica Joossens, Maria Dolores Gómez, Enrico Scarpella, Maria Njo, Yuji Ke, Eline Verhelst and Jim Renema for helpful discussions. We are thankful to the VIB Single Cell Core, VIB Flow Core Ghent, VIB Nucleomics and VIB IRC Histocore for support and access to the instrument park (vib.be/technologies). M.V.B and K.V thank the Research Foundation-Flanders, ELIXIR Belgium (I000323N) for support.

This work was funded by the Ghent University Special Research Fund (BOF20/GOA/012) to T.K., B.D.R. and T.B.; T.E was supported by the European Research Council (ERC) CoG PIPELINES grant (101043257) to B.D.R. This work was further funded by PID2022-138989OB-I00, MCIN/AEI 10.13039/501100011033/ and by FEDER UE. Castilla La Mancha Government (SBPLY/21/180501/000033) to C. E. A.G.R. received an EMBO scientific exchange grant.

## Author contributions

Conceptualization, C.E., T.B., T.K. and B.D.R., data curation, M.S.-S and T.E.; formal analysis, M.S.-S, T.E, M.V.B. and K.V., funding acquisition, C.E., T.B., T.K. and B.D.R., investigation, M.S.-S., A.G.R., M.D., M.M., C.G., T.E., P.A.-U., and R.T.B; methodology, M.S.-S., A.G.R., M.D., M.M., T.E., M.V.B. and K.V., supervision, M.S.-S., C.E., T.B., T.K. and B.D.R.; visualization, M.S.-S, writing-original draft, M.S.-S. and B.D.R. writing – reviewing & editing, M.S.-S., A.G.R., M.D., M.M., C.G., T.E., P.A.-U., M.V.B., R.T.B, K.V., C.E., T.B., T.K. and B.D.R.

## Methods

### Plant material and growth conditions

For time course infection analysis, scRNA-seq and gene expression analysis experiments the following growth conditions were used. Seeds of *Arabidopsis* were surface sterilized with bleach and ethanol and sown on a nylon mesh (20 µm) within square plates containing solid ½ MS without sucrose (1% agar). Seeds were stratified at 4°C for two days and then transferred to long day photoperiod condition (16h/8h) at 22°C. For gene expression analysis using *pWOX5::erGFP* the sterilized seeds were sown in plates containing solid Gamborg B5 medium with vitamins (1.5% sacarose, 0.6% agar). Seeds of *Oryza sativa* spp. *japonica* cultivar Nipponbare were were surface sterilized with 70% ethanol for 1 minute (min), then with bleach (1.5% sodium hypochlorite) for 30 min and then incubated in sterile water for at least 20 min. Seeds were sown on a nylon mesh (20 µm) within square containing solid ½ MS without sucrose (1% agar) and transferred to a long day photoperiod condition (14h/10h) at 24°C. For nematode infection tests the following growth conditions were used. Seeds of *Arabidopsis* were sterilized and growth as previously described^78^ with the following variations: 20-25 seeds were placed on 120mm square petri dishes. Rice seeds were germinated in darkness for 4 days at 30°C. Sprouted seeds were transferred to polyvinyl chloride (PVC) tubes (diameter 3 cm, length 18 cm) containing SAP substrate (sand mixed with Absorbent Polymer AquaPerla; DCM)^79^. They were further grown in a rice growth room to a photoperiod condition (12h/12h) at 28°C (150 μmol·m−2·s−1, and relative humidity of 75%). The *Arabidopsis thaliana* (L.) Heynh. Col-0 ecotype was used as wild-type and background for all the experiments in *Arabidopsis*, while *Oryza sativa* spp. *japonica* cultivar Nipponbare was used in rice. *Arabidopsis* plant lines used in this study were previously described: *pGATA23::NLS-GFP-GUS*^80^, *pPUCHI::GUS*^81^, *pGLV5::NLS-2xGFP*^50^, *pWOX5::erGFP*^82^, *pEXPA4::NLS-GFP-GUS*^83^, *pATHB2::ATHB2-GUS*^84^, *Atathb2-1*^70^. For rice functional characterization, wild-type and *Oshox28* CRISPR mutant seeds (catalog number: CCRM-0025) were obtained from Creative Biogene.

### Nematode maintenance, extraction and surface sterilization

A sterile *Meloidogyne javanica* Treub culture was maintained *in vitro* in *Cucumis sativus* roots as described previously^85^. Infective J2s were obtained after hatching egg masses from the cucumber roots in sterile tap water. A *Meloidogyne graminicola* culture was maintained on susceptible host plants: *Oryza sativa cv. Nipponbare* and Echinocloa cruss-galli. Infective J2s were extracted from infected plants following a modified Baermann funnel method^86^. When needed, J2s were surfaced sterilized with antibiotics (50 µg gentamycin, 100 µg kanamycin, 100 µg spectinomycin) for 30 min and Hospital Antiseptic concentrate (MOLNLYCKE, 3.3 µl/ml) for 5 min, then the J2s were washed with sterile tap water three times.

### Nematode infection experiments

For time course infection analysis, scRNA-seq and gene expression anaylisis experiments the following protocols were used. *Arabidopsis* roots from 5-day-old seedlings were inoculated with 15 J2 nematodes per plant. After nematode inoculation, 23% Pluronic F-127 solution was applied above the root tips. For mock treatment, only 23% Pluronic F-127 solution was applied in the roots of the plants. Plants were transferred then again to long day photoperiod condition and covered with a gauze to reduce light intensity until harvesting. Rice roots from 5-day-old seedlings were inoculated with 50 surface sterilized J2 nematodes per plant. For nematode infection tests the following protocols were used. *Arabidopsis* roots from 5-day-old seedlings were inoculated with 15 J2 nematodes per plant. After inoculation plants were growth in darkness for 4 days. The plates were then placed at 45 degrees and covered with a gauze to protect from excess light till the end of the experiment. Galls and egg masses in the main root were counted at 10 dpi and 45 dpi respectively under a stereomicroscope (Leica MZ6; Leica, Wetzlar, Germany). Two replicates were performed for both infection and reproduction test with a total of at least 300 plants for the infection test and 200 plants for the reproduction test. Rice roots from two-week-old plants were inoculated with 1000 J2s per plant. Plant susceptibility was assessed at 7 dpi by counting galls and total nematodes using the acid fuchsin staining technique. Galls and nematodes were counted under a stereomicroscope (SMZ1270, Nikon). The infection experiment was repeated twice, each time using 13-15 plants per genotype.

### GUS activity and histological analysis

For GUS assays, seedlings were incubated in 90% acetone for 20 min in ice. Then, they were washed twice in 0.1 M phosphate buffer (pH 7.2) and incubated for 1h to overnight in the GUS incubation buffer (1 mg/ml 5-bromo-4-chloro-3-indolyl β-D-glucuronic acid, 0.5 mM potassium ferrocyanide, and 0.5 mM potassium ferricyanide in 0.1 M phosphate buffer). The reaction was stopped by washing three times in 0.1 M phosphate buffer and the samples were subsequently processed for nematode staining. For nematode staining, rice infected root tips or galls were cut with a scalpel and treated with bleach (4%) for 10 min, washed in tap water for 15 min, stained in boiling acid fuchsin solution (0.013% acid fuchsin in 25% acetic acid) for 3 min, washed twice with tap water and then cleared with de-staining solution (33% glycerol, 33% lactic acid). The same procedure was used for Arabidopsis infected roots with some changes (1% bleach and 0.039% acid fuchsin). Photographs were taken with either a stereomicroscope (Leica M165 FC or Olympus SZX16) or with a light microscope (Olympus Bx51). When needed, samples were washed in 0.1 M sodium phosphate buffer and processed for Technovit embedding. For detailed visualization of roots, root tips or galls were fixed in 2% (w/v) paraformaldehyde and 1% (v/v) glutaraldehyde in sodium phosphate buffer (0.1 M NaH2PO4.H2O, pH 7.2) overnight at 4 °C. Then the tissue was dehydrated and embedded in Technovit 7100 (Heraeus Kulzer, Hanau, Germany) according to the manufacturer’s protocol. Sections of 4 µm were cut with a microtome (Leica Reichert-Jung 2050) and mounted on SuperFrost® Plus slides (Menzel-Glaser). The sections were stained with 0.1% Toluidine blue for 1 min or 0.05% Ruthenium red for 8 min and briefly rinsed in distilled water. After drying, the sections were mounted in DPX mounting medium (Sigma Aldrich, 06522).

### Protoplast isolation and FACS

*Arabidopsis* root tips (approx. 1 cm) were harvested at 36 hpi and incubated in Solution B [1.5% (wt/vol) Cellulase YC and 0.1% (wt/vol) pectolyase in Solution A (400 mM mannitol, 20 mM MES, 20 mM KCl, 10 mM CaCl_2_, pH 5.7)] for approximately 1 h at room temperature in darkness. Cells were filtered through a 70 µM cell strainer and spin down at 200 x g for 6 min and resuspended in Solution A. Cells were sorted based on size and granularity parameters using a BD Biosciences FACS Discover S8 enabled prototype cell sorter.

Rice root tips (approx. 0.5 cm) were harvested and incubated in protoplast solution for 2 h applying vacuum for the first 15 min. The protoplast solution was prepared by preparing the enzymatic solution (400 mM mannitol, 20 mM MES, 20 mM KCl, 1.25% Cellulase RS, 1.25% Cellulase R10, 0.3% Macerozyme R10, 0.12% Pectolyase, pH 5.7), heating it up for 10 min in a 60°C water bath and then cooling it to room temperature to add the rest of the compounds (10 mM CaCl2, 0.1% BSA, 0.018% (v/v) β-mercaptoethanol). After incubation, the solution was pipetted up and down thoroughly with a 1 ml tip until it got cloudy for releasing the protoplasts^2^. Cells were filtered through a 70 µM cell strainer and spin down at 200 x g for 6 min and resuspended in 8% mannitol twice. Then, cells were filtered through a 40 µM cell.

### 10X Genomics sample preparation, library construction and sequencing

For the rice samples, the concentration of filtered protoplasts was determined and around 16k protoplasts were loaded on a Chromium GemCode Single Cell Instrument (10x Genomics) using Chromium Next GEM Single Cell 3ʹ Reagent Kit v3.1 (10x Genomics) according to the manufacturer’s instructions. The first two replicates were processed by subdividing the same batch of protoplasted tissue in two reactions that were loaded independently. For the Arabidopsis samples, 100k sorted protoplasts were centrifuged at 4°C for 6 min at 400 g, resuspended in washing solution and loaded on a Chromium GemCode Single Cell Instrument (10x Genomics) using Chromium Next GEM Single Cell 3ʹ Reagent Kit v3.1 (10x Genomics) according to the manufacturer’s instructions. Sequencing libraries were loaded on an Illumina NovaSeq6000 flow cell and sequenced following recommendations of 10x Genomics at the VIB Nucleomics core (VIB, Leuven).

### Raw data processing and generation of gene expression matrix

The FASTQ files obtained after demultiplexing were used as input for “cellranger count” (version 6.0.0, 10x Genomics) to map the reads to the respective reference genomes, being TAIR10.40 for Arabidopsis and IRGSP.49 for rice. Preprocessing of the data was done by the scater^87^ (v1.20.1) R package according to the workflow proposed by the Marioni lab^88^. Outlier cells in rice were defined as having fewer than 1000 expressed genes or 1200 UMIs or having more than 10% mitochondrial or 15% chloroplast transcripts. In Arabidopsis, outlier cells had fewer than 800 expressed genes or 1200 UMIs, or more than 15% chloroplast or mitochondrial transcripts.

### Data analysis (clustering, filtering, replicate/batch, subsampling clustering, degs clusters)

The data analysis of scRNA-seq was performed with Seurat (v4)^89^ using R^90^ and RStudio. For each species, all the filtered datasets from infected and mock samples were merged and normalized with SCTransform function using the 3000 variable features. For dimensionality reduction, a principal component analysis (PCA) was performed using the function RunPCA followed by a non-linear dimensional reduction with the function RunUMAP using the top 30 principal components. For grouping cells within clusters, the FindNeighbors function was applied using the top 30 principal components from the PCA reduction and then the FindClusters function with a resolution of 0.8 for *Arabidopsis* dataset and 0.6 for the rice dataset. For identifying marker genes for every cluster, the function FindAllMarkers was used with default parameters.

### Cell type and cell cycle phase annotation

For annotating cells depending on the cell type in rice, datasets from mock samples (R1, R2 and R3; **Supplementary Table 1**) were merged and normalized using the 3000 top variable features and dimensionality reduction and clustering was performed as described before. We checked the gene expression from cell type specific markers experimentally validated by RNA *in situ* hybridization in this paper or previously somewhere else (**Supplementary Table 3**). Based on this information, clusters were annotated manually. This dataset was used then as a reference for transferring the cell type annotation to the complete merged dataset. Briefly, the FindTransferAnchors function using the 30 top principal components from the PCA reduction was used with the annotated mock dataset as reference and the complete merged dataset as query. The predicted cell type with higher score was assigned for every individual cell with the TransferData function. For annotating cells depending on the cell type in *Arabidopsis*, a previously root scRNA-seq atlas was used as reference^49^. For every individual sample dataset before merging the FindTransferAnchors function using the 30 top principal components from the PCA reduction was applied with the published dataset as reference. The predicted cell type with higher score was assigned for every individual cell with the TransferData function. For annotating cells depending on the cell cycle phase the function CellCycleScoring from Seurat was used in every individual sample dataset before merging. The genes used as markers of the S phase and the G2/M phase are listed in **Supplementary Table 3**. For rice, orthologous genes of *Arabidopsis* known cell cycle genes markers were used. For *Arabidopsis*, the top differentially expressed genes for the clusters grouping dividing cells corresponding to the S phase and the G2/M phase from a previous root single cell dataset were used^49^.

### Subsampling and cluster enrichment analysis

To mitigate potential bias in the enrichment analysis of cells from infected samples during clustering, due to unequal cell numbers between treatment conditions, a subsampling strategy combined with cluster analysis was employed for both *Arabidopsis* and rice datasets. Each dataset was subsampled 10 times with replacement. In each iteration, 10,000 cells from the mock treatment and 10,000 cells from the infected treatment were randomly selected, and the standard pipeline for unsupervised clustering, as described above, was applied. For each iteration, the first clustering resolution that identified more than 14 communities was selected to ensure a consistent number of clusters for downstream analysis. Then, the frequency of infected cells within each cluster was calculated, and the clusters were ranked according to this value. To assess the stability of infection-associated populations, the frequency distribution of cells from the original “nematode clusters” was evaluated across the new clusters in each subsample.

### Visualization gene expression (feature plot/dotplot)

Visualization of gene expression on the UMAP projection was performed with the function FeaturePlot from Seurat using the log-normalized counts from the SCTransform assay. Visualization of average gene expression across clusters was performed with the function DotPlot from Seurat using the log-normalized counts from the SCTransform assay.

### Nematode score assignment

Upregulated genes in roots infected with RKN compared to non-infected roots were retrieved from transcriptome data analysis from previous studies^27,30,31,91–94^ (Details in **Supplementary Table 1**). Five studies were included for each species and genes upregulated in at least three of them were taken in account to constitute the “nematode core gene set” (288 genes in Arabidopsis and 109 genes in rice) (**Supplementary Table 1**). The normalized count data for these genes was extracted using the SCT normalized counts. The expression values were then z-score normalized across all cells for every gene. Then, the nematode score for every cell was calculated by summing the z-scores and rescaled [-1, 5]. To classify cells as positive or negative for the nematode response, a threshold score was established for each species (0.5 for Arabidopsis and 1 for rice dataset) by minimizing the percentage of cells from the mock sample (less than 2%) that exceeded the score, thus optimizing for specificity of the nematode response.

### Gene orthology inference

In order to retrieve orthogroups and genes with one-to-one ortholog relationship between rice and *Arabidopsis*, a custom PLAZA^95^ instance was generated with 8 different monocots and dicots plant species, as well as *Amborella trichopoda* as outgroup species (**Supplementary Table 2**). The annotation data for the custom PLAZA instance was retrieved from PLAZA 5.0. The build procedure of this custom PLAZA instance is an updated version of the PLAZA 5.0 build pipeline. Sequence similarity scores between all protein coding genes were generated using DIAMOND v0.9.30^96^. Subsequently, gene families were created using TRIBE-MCL v1.0^97^ and OrthoFinder v2.5.3^98^. Phylogenetic trees for both homologous and orthologous gene families were constructed with FastTree v2.1.10^99^, after the generation of protein multiple sequence alignments with MAFFT v7.453^100^. The reconciliation of the phylogenetic trees with the species tree was performed using NOTUNG v2.6^101^. All programs were executed as part of a Nextflow pipeline^102^. For the identification of the one-to-one orthologous genes, an integrative approach was used. For every gene in *Arabidopsis* the orthologs in rice were inferred by three types of analysis: best-BLAST hit approach, OrthoFinder gene families, or gene family phylogenetic trees. Only those gene pairs supported by at least two analyses were retained as orthologs. These results were further filtered by retaining only the orthologs for which the number of analysis types was the highest per query gene. These orthologous relationships were also evaluated in the other direction (rice to *Arabidopsis*). Only when the same rice-*Arabidopsis* gene pair met these requirements for both directions was it considered as the one-to-one ortholog gene pair.

### Cross-species integration datasets

First, the raw matrixes for each species were filtered to retain only the genes with one-to-one orthologs between rice and Arabidopsis. The gene identifiers from rice were translated to the corresponding *Arabidopsis* gene identifiers. Each dataset was normalized with the SCTransform function from Seurat with default parameters. Then, 3000 features for integrating the datasets were selected applying the function SelectIntegationFeatures from Seurat. Data was prepared with the function PrepSCTIntegration, followed by the function FindIntegrationAnchors with the normalization.method = “SCT” and using Canonical correlation analysis as dimensional reduction method to find the anchors. Lastly, the data was integrated with the IntegrateData function using the anchors previously identified as anchorset. The integrated dataset was then processed according to the standard pipeline. The function RunPCA was used using the first 50 principal components followed by a non-linear dimensional reduction with the function RunUMAP using the top 40 principal components. The function FindNeighbours using the pca as reduction and the first 40 PCs and the FindClusters with resolution 0.4 to obtaining the clusters of the integrated dataset.

### Cross-species pairwise cluster correlations

For each species the average gene expression per cluster was calculated using the function from Seurat AverageExpression and using the SCT normalized data. This resulted in a matrix for each species where the rows represent the genes and the columns the clusters, with the values representing the aggregated gene expression of all the cells within a cluster. Then, two different approaches were applied to calculate pairwise cluster correlations between the clusters of rice and *Arabidopsis* datasets.

#### Correlation between clusters using gene specificity index values

Each aggregated gene expression matrix was filtered to retain only the pair of genes one-to-one ortholog that are expressed in both datasets, and we translated the identifiers of the rice genes into the Arabidopsis identifiers. Expression values were converted to gene specificity index^43^. Briefly, the expression of a gene within a cluster is divided by the mean expression of the gene across all the clusters to obtain the gene specificity index. Pairwise cluster correlations were computed using the Spearman correlation. The p-value of the correlation was calculated through a permutation test with 1000 iterations. In every iteration the rows within each matrix were randomly shuffled while maintaining the internal structure. The Spearman correlation was calculated between these randomized matrices. The p-value was determined by comparing the absolute value of the original correlation with the absolute value of the permuted correlations. The p-value represents the proportion of cases where the permuted correlations exceeded the original correlation. Only those pairwise correlations with a p-value < 0.05 were considered statistically significant.

#### Correlation between clusters using scores of biological process GO terms enrichment

In the second strategy, the aggregated gene expression matrices were used to perform ssGSEA. This allowed us to obtain a new matrix that every row represents a gene set, in this case a biological process (BP) GO term, every column represents a cluster, and the values represent the Normalized Enrichment Scores (NES) obtained from the analysis. First, all the children GO terms of the term BP (GO:0008150) were selected (https://release.geneontology.org, GO release 2021-09-01). For each species the gene sets corresponding to every BP GO term were retrieve from PLAZA monocots 5.0. Only the genes that are present in the aggregated gene expression matrix were retained. Then, only GO terms that contain between 15 and 500 genes and that were common for both species were kept. The gene expression matrices were converted to the BP GO terms normalized enrichment scores by applying the function ssGSEA from the package corto^103^. Pairwise cluster correlations with the transformed matrices were computed with the Spearman correlation and the p-value was calculated as above.

### Analysis variability one-to-one ortholog (subset)

To evaluate whether the use of the subset one-to-one orthologous genes may bias the cluster cross-species comparison, ssGSEA was performed within the marker genes of every cluster in the *Arabidopsis* and rice datasets. Briefly, a ssGSEA was performed on the marker gene (avgFC > 0) list obtained by cluster with the FindAllMarker function from Seurat. The NES and p-values of the on-to-one ortholog gene set and ten random gene sets with the same size were obtained for every cluster with the function ssGSEA from the package corto.

### Single-cell gene regulatory networks using MINI-EX

For identifying single cell gene regulatory networks MINI-EX v1 was used^52^. As MINI-EX does not implement an approach to handle a control/treatment set up, we developed a custom strategy for this purpose. The cells were annotated depending on the cell type annotation and the treatment (e.g, “CortexInfected”). Marker genes were identified for the clusters with the new annotation with the function FindAllMarkers from Seurat and only positive markers were kept. For the rice dataset, RAP identifiers were translated to MSU identifiers to be compatible with the annotation implemented in MINI-EX. The transcription factor lists and binding sites files available in MINI-EX for *Arabidopsis* and rice were used. Finally, MINI-EX pipeline was run using the top 200 marker genes and taking in account TFs that were expressed at least in 8% of the cells within a cluster.

### Dioxigenin-labeled RNA probe synthesis

RNA probes were designed to hybridize specifically with the complete transcript sequence of interest and avoiding including genome repetitive regions. Specific oligonucleotides (**Supplementary Table 6**) were used to amplify the region of interest from cDNA of rice roots. The PCR product was ligated into the pJET blunt 1.2 vector following the ConeJET PCR cloning kit (Thermo Fisher) instructions. The plasmid containing the antisense fragment was selected and the fragment containing the T7 initiator was amplified using the corresponding oligonucleotides (**Supplementary Table 6**). The PCR product was concentrated using DNA Clean & Concentrator-5 (Zymo Research). The transcription reaction was performed using the MEGAscript™ T7 Transcription Kit (Invitrogen). Briefly, a mix of nucleotides was prepared using ATP, CTP and GTP from the kit and the labeled UTP was prepared mixing with Dioxigenin-11-UTP (Roche):UTP in a 2:1 ratio. Then, 1 µg from the DNA template was incubated with every nucleotide at a final concentration of 12.16 nM and 1.5 µl of the enzyme mix in a final volume of 15 µl. The reaction was incubated overnight at 37°C and the DNA template was removed by adding 1 µl of turbo DNase I from the kit at 37°C for 20 min. The volume was brought up to 100 µl by adding RNAse-free water. Probes were hydrolyzed by adding an equal volume of 2x Carbonate Hydrolysis buffer (80 mM NaHCO_3_, 120 mM Na_2_CO_3_, prepared in RNAse-free water), neutralized by adding 10 µl of 10% acetic acid and precipitated by adding 40 µl of LiCl (4M) followed by 400 µl of 100% ethanol. The hydrolyzed probes were washed twice with 70% ethanol and finally resuspended in 100 µl 50% de-ionized formamide. The DIG-labeled hydrolyzed probes were stored at - 80°C.

### RNA *in situ* hybridization (ISH) in rice

#### RNA ISH in paraffin sections with dioxigenin-labeled probes

A protocol optimized for monocots root tips previously described was used with some modifications^104^. For all the pertinent solutions, water treated with diethyl pyrocarbonate (0.1% v/v, DEPC) was used. Root tips were harvested and incubated in fixative solution FAA (3.7% formaldehyde, 5% glacial acetic acid, 50% ethanol in DEPC water) overnight at 4°C. Samples were dehydrated in a series of washes at room temperature for at least 30 min in every step: PBS (0.13 M NaCl, 0.007 M Na_2_HPO_4_, 0.003 M NaH_2_PO_4_.H_2_O, in DEPC water), 50% ethanol, 70% ethanol, 85% ethanol, 95% ethanol, 1x ethanol. Then, samples were washed again in 100% ethanol overnight at 4°C. Next day, samples were washed at room temperature for 2 h in 100% ethanol and permeabilized in a series of washes at room temperature for 1 h in every step: 25% HistoClear-II:75%ethanol, 50%HistoClear-II:50%ethanol, 75% HistoClear-II:25% ethanol, 100% HistoClear-II (three times). Then, Paraplast Plus® (Sigma-Aldrich) chips were added to approximately half the volume of the HistoClear-II and samples were kept there overnight. The samples were incubated at 60°C until the paraffin was dissolved. The liquid was replaced with freshly molten paraffin and kept overnight at 60°C. During the next two days, the paraffin was replaced every morning and evening with fresh melted paraffin. Paraffin molds were prepared in a Epredia™ HistoStar™ Embedding Workstation. Sections of 10 µm were cut with a microtome (Leica Reichert-Jung 2040) and mounted on SuperFrost® Plus slides (Menzel-Glaser). Sections on the slides were de-paraffined and rehydrated in a series of washes at room temperature: 100% HistoClear-II (twice) for 10 min, 100% EtOH (twice), 95% ethanol, 70% ethanol, 50% ethanol, 30% ethanol and DEPC water, for 30 seconds (s) each, and 1x DPBS for two min. Sections were permeabilized by incubating 10 min in pre-warmed pronase buffer at 37°C (50 mM This-HCl pH 7.5, 5 mM EDTA pH 8, 0.12 mg/ml Protease (Sigma-Aldrich) in 1x PBS), followed by an incubation in 0.2% glycine (in 1x DPBS) for 2 min and washing again in 1xPBS for 2 min. Samples were re-fixed by incubating in 3.7% formaldehyde (in DEPC water) for 10 min and washed in 1x DPBS for 2 min. Samples were acetylated by incubating in 0.1 M Triethanolamine (Sigma-Aldrich) and 5 µl/ml of acetic acid (in DEPC water) for 10 min and washed in 1x PBS for 2 min. Sections were then dehydrated in a series of washes: 30% ethanol, 50% ethanol, 70 % ethanol, 95% ethanol, 30 s each, 100% ethanol for 2 min (twice) and then kept in a humid environment with 100% ethanol at 4°C until they are used in the next step. The hydrolyzed probes were diluted in 50% formamide (2 µl probe stock in 98 µl) and denatured at 95°C for 3 min, then placed on ice for 1 min and back to room temperature. Then, 400 µl of the hybridization buffer (0.3 M NaCl, 0.01M Tris-Cl pH 6.8, 0.01 M NaPO_4_, 5 mM EDTA, 50% de-ionized formamide, 12.5% Dextran sulfate, 1.25 mg/ml tRNA, 1 mL 1.25x Denhardt’s solution) were added to the probes. The slides were hybridized with 125 µl of the probes in the buffer at 60°C overnight in a humid chamber. Next day, probes were removed by washing in 0.2x SSC for 1 h at 55°C (three times). Slides were rinsed in 1x PBS for 5 min at room temperature and then they were incubated in blocking solution I (10 mg/ml Boehringer Block reagent, 0.15 M NaCl, 0.1 M Tris-HCl pH 7.5) for 45 min. The solution was then replaced by the blocking solution II (10 mg/ml BSA, 0.1 M Tris-HCl pH 7.5, 0.15 M NaCl, 0,003% Triton X-100 (v/v)) and incubated for 45 min. Then, slides were incubated with the anti-Dig alkaline phosphatase conjugated antibody (0.8 µl/ml in blocking solution II) at room temperature for 2 h. The antibody solution was replaced by blocking solution II and was for 20 min (four times). The solution was replaced by freshly made Buffer C (100 mM Tris-HCl pH 9.8, 50 mM MgCl_2_, 100 mM NaCl) for 15 min. Lastly, slides were incubated in NBT/BCIP (20 µl/ml in Buffer C) in darkness at room temperature overnight. Slides wre rinsed in water for 3 min to stop the reaction and dehydrate rapidaly in an alcohol series before mounting with Permount (Thermofisher) with a cover slide. Samples were imaged with a light microscope (Olympus Bx51).

#### HCR RNA fluorescence in situ hybridization

For half-mount fluorescent in situ hybridization (FISH), FISH was performed following a protocol optimized for monocot root tissue as described previously^44^. Gene specific probes (hairpin chain reaction (HCR) RNA-FISH) and reagents (probe hybridization buffer, probe wash buffer and amplification buffer) were obtained from Molecular Instruments. For the rest of the solutions, DEPC water was used. Gene probes and fluorescently labeled hairpins used are described in Supplementary Table 6. Root tips from infected and mock roots were gently cut superficially and longitudinally with a 30° microscalpel previously dipped with in a fixative solution FAA. Root tips (2 cm.) were then harvested and transferred to FAA under vacuum for 1 h and then for 45 min in an orbital shaker at room temperature. Samples were dehydrated at room temperature: 15 min in 70% ethanol, 15 min in 90% ethanol, 15 min in 100% ethanol (twice), 15 min in 100% methanol (twice), 15 min in 100% ethanol (twice). Then, samples were permeabilized by incubating 30 min in 50%Histo-ClearII:50% ethanol and 30 min in 1000% Histo-ClearII (twice). Every time, vacuum was applied for the first 10 min. Samples were rehydrated in a series of washes: 15 min in 50%Histo-ClearII:50% ethanol, 15 min in 100% ethanol, 15 min in 50% ethanol:50% DPBS-T (0.1% Tween-20, 1x PBS in DEPC water), 15 min in DPBS-T (twice). Samples were then digested in Proteinase K buffer (0.1 M Tris HCl (pH 8), 0.05 M EDTA (pH 8), Proteinase K 80 μg/ml) for 5 min at room temperature under vacuum and then for 25 min at 37°C. Then, samples were washed for 15 min DPBS-T (twice) and refixed for 40 min in fixative solution II (3.7% formaldehyde in DPBS-T), applying vacuum for the first 10 min. Afterwards, samples were washed again for 15 min in DPBS-T (twice). For probe hybridization, samples were incubated for 10 min under vacuum in HCR probe hybridization buffer and then for 1 h at 37°C. The buffer was replaced with HCR probe hybridization buffer containing the corresponding gene probes (1-5 μM each) and incubated overnight (20 h) at 37°C. Both infected and mock roots were incubated at the same probe concentration. Samples were washed for 30 min in HCR Probe Wash buffer at 37°C (twice) and then for 5 min in 5x SSC-T (0.75 M NaCl, 0.075M sodium citrate, 0.1% Tween-20 in DEPC water, pH = 7) at room temperature. Then, samples were transferred to the amplification buffer for 10 min under vacuum and for 50 min at room temperature. In the meantime, each fluorescently labeled hairpin was prepared by snap cooling the required volume of the 3 μM stock in hairpin storage buffer by heating at 95°C for 90s in thermocycler and cooling to room temperature in the dark for 30 min. Amplification solution was prepared by combining snap-cooled h1 and h2 hairpins in HCR amplification buffer (0.06 μM of every hairpin). Samples were transferred to the amplification buffer containing the hairpins and incubated overnight in darkness at room temperature. Samples were washed for 20 min in 5x SSC-T (three times) and transferred to ClearSee (reference, recipe) for several days. For visualization, samples were embedded in 3% agarose (in 1x DPBS). Transversal sections of the roots were obtained by cutting the agarose blocks into 200 μm sections using a vibratome (Leica VT1200 S). Then sections were incubated in Calcofluor (0.01% in ClearSee) directly in the slide previously to imaging.

### Imaging

Confocal microscopy of *Arabidopsis* marker lines observation was performed using a Leica TCS SP8 spectral microscope. First, harvested root tips were fixed in 4% PFA for 30 min under vacuum, then washed in PBS and clarified in ClearSee at least overnight. Cell walls were stained overnight with Direct Red 23 (0.01% in ClearSee) and nucleic acids were stained with SYTOX^TM^ ORANGE (5µM). Green fluorescent protein, Direct Red 23 and SYTOX Orange were excited with lasers at 488 nm, 552 nm and 552 nm; and Hybrid detector was set at 500-530nm, 580-625 nm respectively. Confocal microscopy of FISH in rice root sections was performed using a Leica TCS SP8 spectral microscope. Calcofluor, orthogonal amplifiers 488, 546 and 647 were excited with lasers at 405 nm, 488 nm, 560 nm and 647 nm and detected at the ranges of 430-486 nm, 497-523 nm, 559-596 nm and 654-707, respectively.

### RNA isolation, library preparation and sequencing

RNA was extracted from rice root tips or protoplasts from rice root tips with the RNeasy Plant Mini Kit (Qiagen) following the manufacturer’s protocol, with an additional sonication step for 30 seconds after addition of buffer RLT. Libraries were prepared using the Quantseq 3′ mRNA prep kit (Lexogen) following the manufacturer’s instructions. After confirming the quality of the libraries with an Agilent Bioanalyzer 2100, the libraries were sequenced on Illumina NextSeq 500, generating single-end 76 bp reads.

### RNA-seq data analysis

First, adapters and low-quality bases were trimmed from the raw reads using Trimmomatic v0.38^105^ with default settings, requiring a minimum length of 20 nt. FastQC v0.11.8 quality control was performed before and after trimming. Trimmed reads were subsequently mapped to the *O. sativa* spp. *japonica* reference genome (IRGSP-1.0/MSU7) with Bowtie2 v2.3.4.3^106^ in end-to-end mode. Count tables were produced on IRGSP-1.0.42 annotation using the SummarizeOverlaps function from the GenomicAlignments package^107^ in R Bioconductor. Differential expression analysis was carried out with DESeq2 v.1..24.0^108^. Genes with fold-change > 1.5 and adjusted p-value < 0.001 were selected as up-regulated by protoplast isolation.

## Extended Data Figure legends

**Extended Data Fig. 1.**
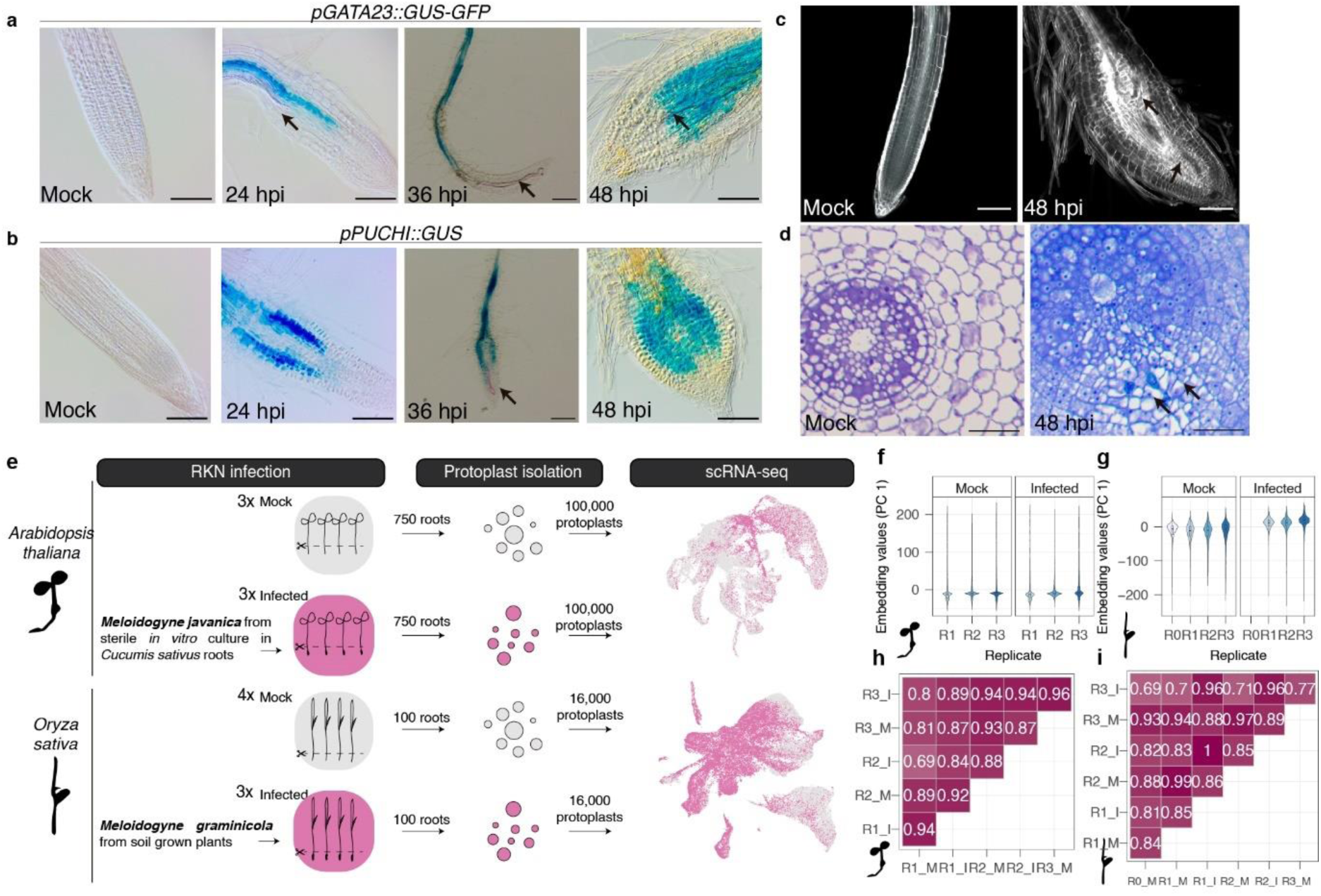
Set up and metrics of *Arabidopsis* and rice scRNA-seq data. **a,b,** Representative *Arabidopsis* roots showing the expression of *GATA23* (**a**) and *PUCHI* (**b**) at different timepoints during *Meloidogyne javanica* infection, as monitored by histochemical detection of GUS (blue substrate). Arrows points towards visible RKN within the roots. Scale bars, 100 μm. **c,** Confocal micrographs of *Arabidopsis* mock roots (left) or infected roots at 48 hpi (right). Cell walls were stained with Direct Red 23 and represented in grey scale. Arrows points towards RKNs. Scale bars, 100 μm. **d,** Left, cross-section of 7-day-old rice mock root. Right, Cross-section of rice infected root at 48 hpi. Arrows points towards RKNs. Scale bars, 100 μm. **e,** Schematic diagram of scRNA-seq experiments. **f,g,** Violin plots of embedding values in the first principal component (PC1) of PCA reduction per cell within the replicates and treatment in *Arabidopsis* (**f**) and rice (**g**) datasets. **h,i,** Pearson correlation coefficient of the average gene expression among replicates in *Arabidiposis* (**h**) and rice (**i**) datasets.

**Extended Data Fig. 2.**
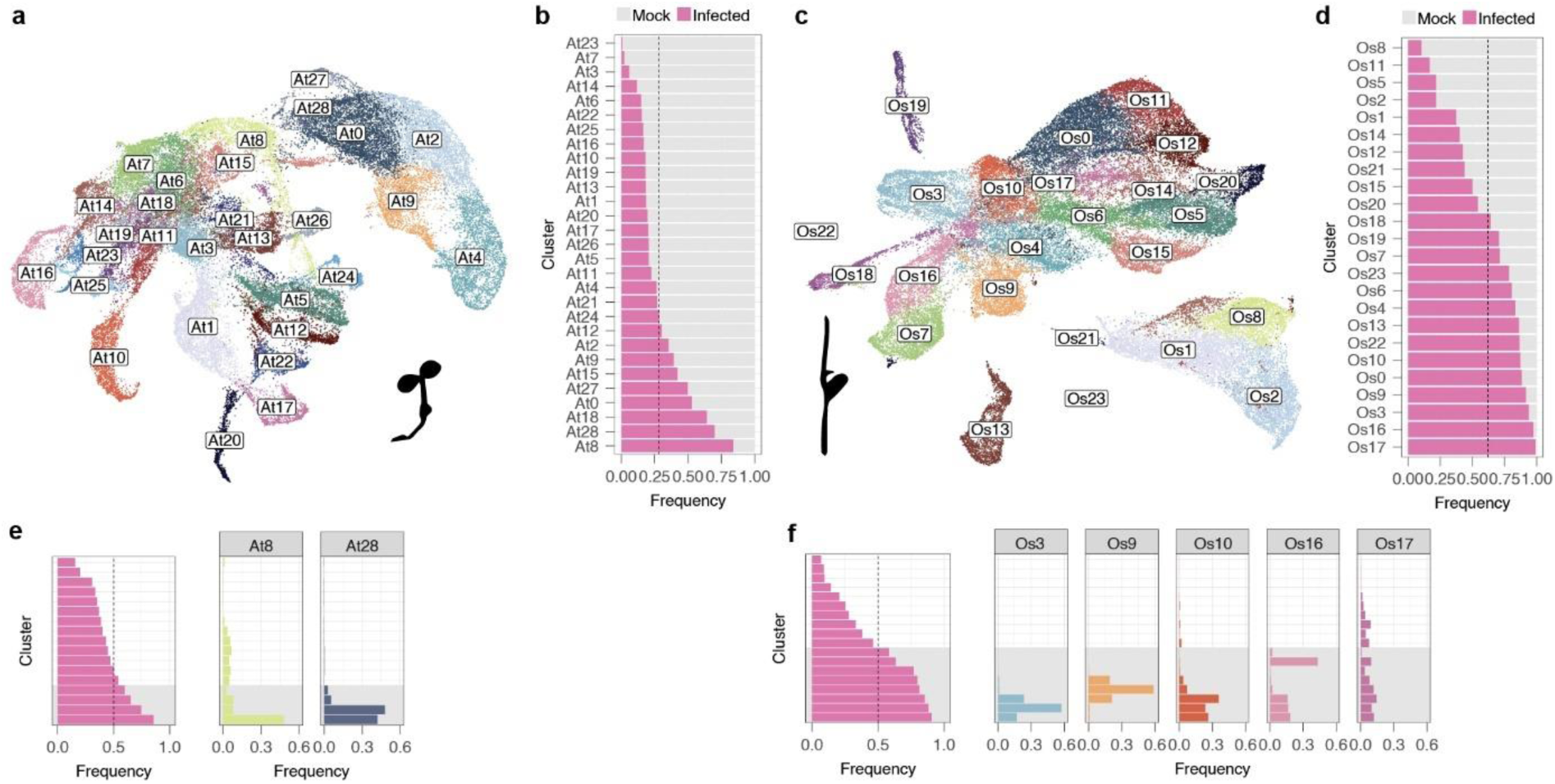
Clustering analysis of *Arabidopsis* and rice datasets. **a,** UMAP representation of the *Arabidopsis* scRNA-seq dataset showing the unsupervised clustering using the Louvain algorithm at a resolution of 0.8. Colors and labels represent distinct clusters. **b,** Frequency of cells from mock or infected samples in the clusters shown in **a**. Dotted line indicates the expected frequency of infected cells. **c,** UMAP representation of the rice scRNA-seq dataset showing the unsupervised clustering using the Louvain algorithm at a resolution of 0.6. Colors and labels represent distinct clusters. **d,** Frequency of cells from mock or infected samples in the clusters shown in **c**. Dotted line indicates the expected frequency of infected cells. **e,f,** Evaluation of consistency in cluster enrichment using balanced datasets by subsampling the *Arabidopsis* (**e**) and rice (**f**) datasets. Left panels, average frequency of cells from infected samples across clusters in subsampled datasets. Bars represent the average frequency of infected cells in each cluster across 10 subsampling iterations, ordered from lowest to highest frequency. The dashed line represents equal distribution between infected and mock cells. Clusters with grey background show higher frequency than 0.5. Right, distribution of cells that belong to the clusters enriched in infected cells in the complete datasets across the new clusters in the subsampled dataset. Bars show the mean frequency of cells from each original selected cluster appearing in the new clusters (ordered as on the left panel) across 10 subsampling iterations.

**Extended Data Fig. 3.**
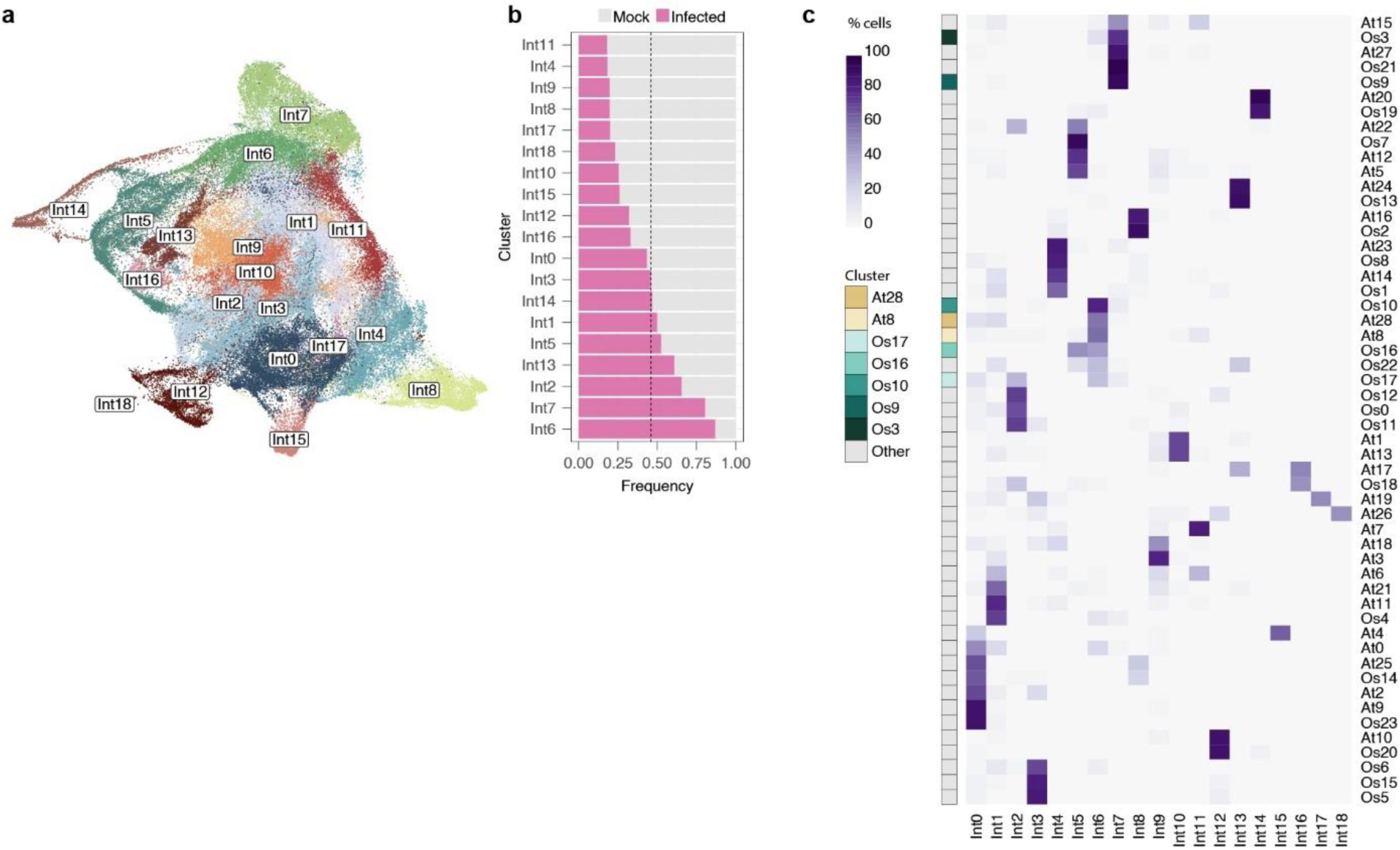
Integration of *Arabidopsis* and rice datasets. **a,** UMAP representation of *Arabidopsis* and rice integrated dataset showing the unsupervised clustering using the Louvain algorithm at a resolution of 0.4. Colors and labels represent distinct clusters. **b,** Frequency of cells from mock or infected samples in the clusters shown in **a**. Dotted line indicates the expected frequency of infected cells. **c,** Heatmap showing the percentage of cells originally assigned to each cluster in the separate *Arabidopsis* and rice datasets that are now distributed across clusters in the integrated dataset shown in **a**.

**Extended Data Fig. 4.**
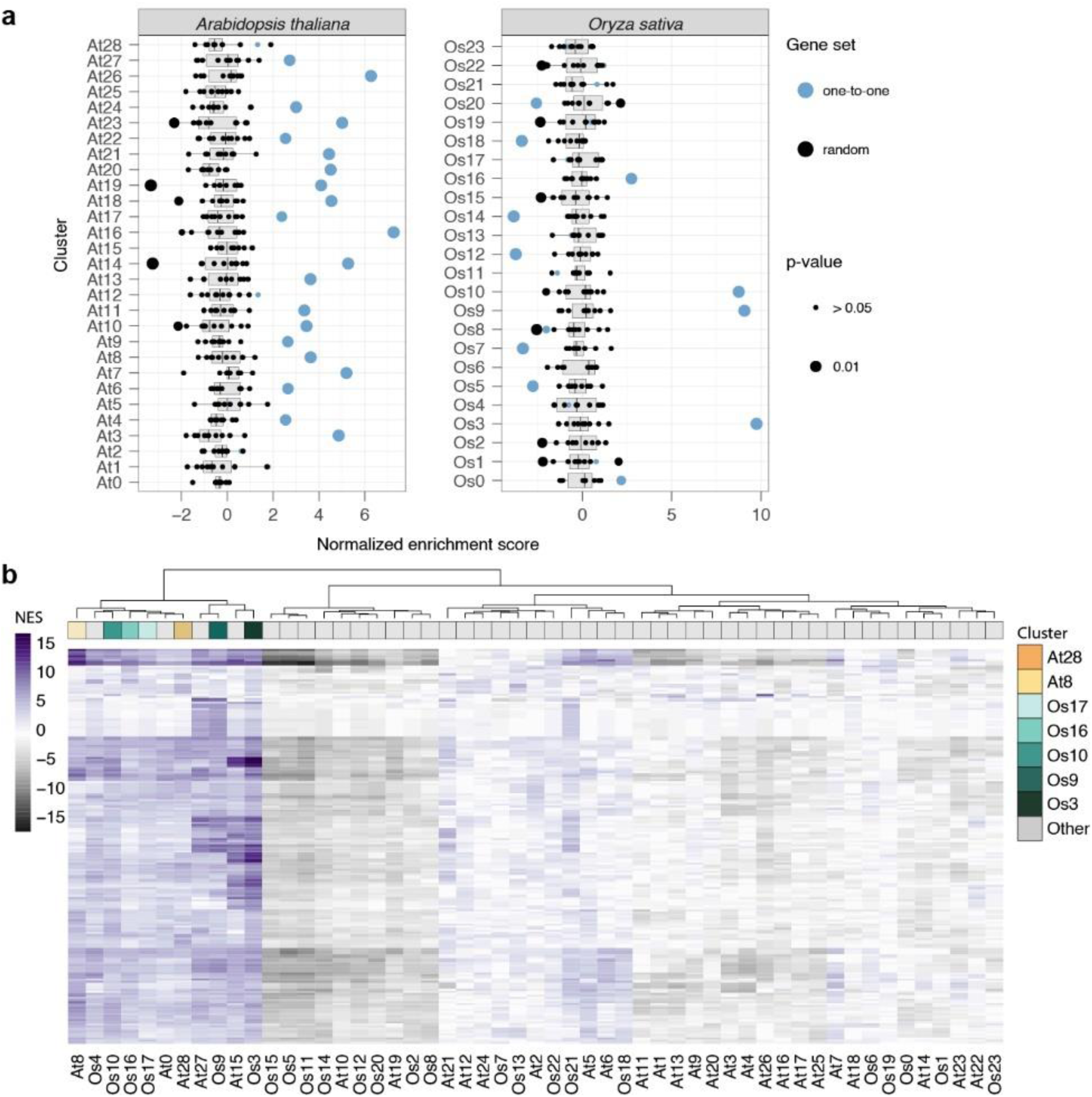
Evaluation of enrichment of one-to-one ortholog and BP GO terms sets. **a,** Enrichment analysis of one-to-one orthologous genes in differentially expressed genes (DEGs) across the *Arabidopsis* (left) and rice (right) clusters. For every cluster, the normalized enrichment scores (NES) are represented for the subset of one-to-one orthologous genes and ten subsets of the same size containing random genes. Big dots represent a significant enrichment (p-value < 0.01). **b,** Heatmap showing the normalized enrichment scores (NES) of the combined top 60 biological process terms that define the “nematode clusters” across *Arabidopsis* and rice clusters. Clusters are grouped using hierarchical clustering based on the distance between their enrichment profiles.

**Extended Data Fig. 5.**
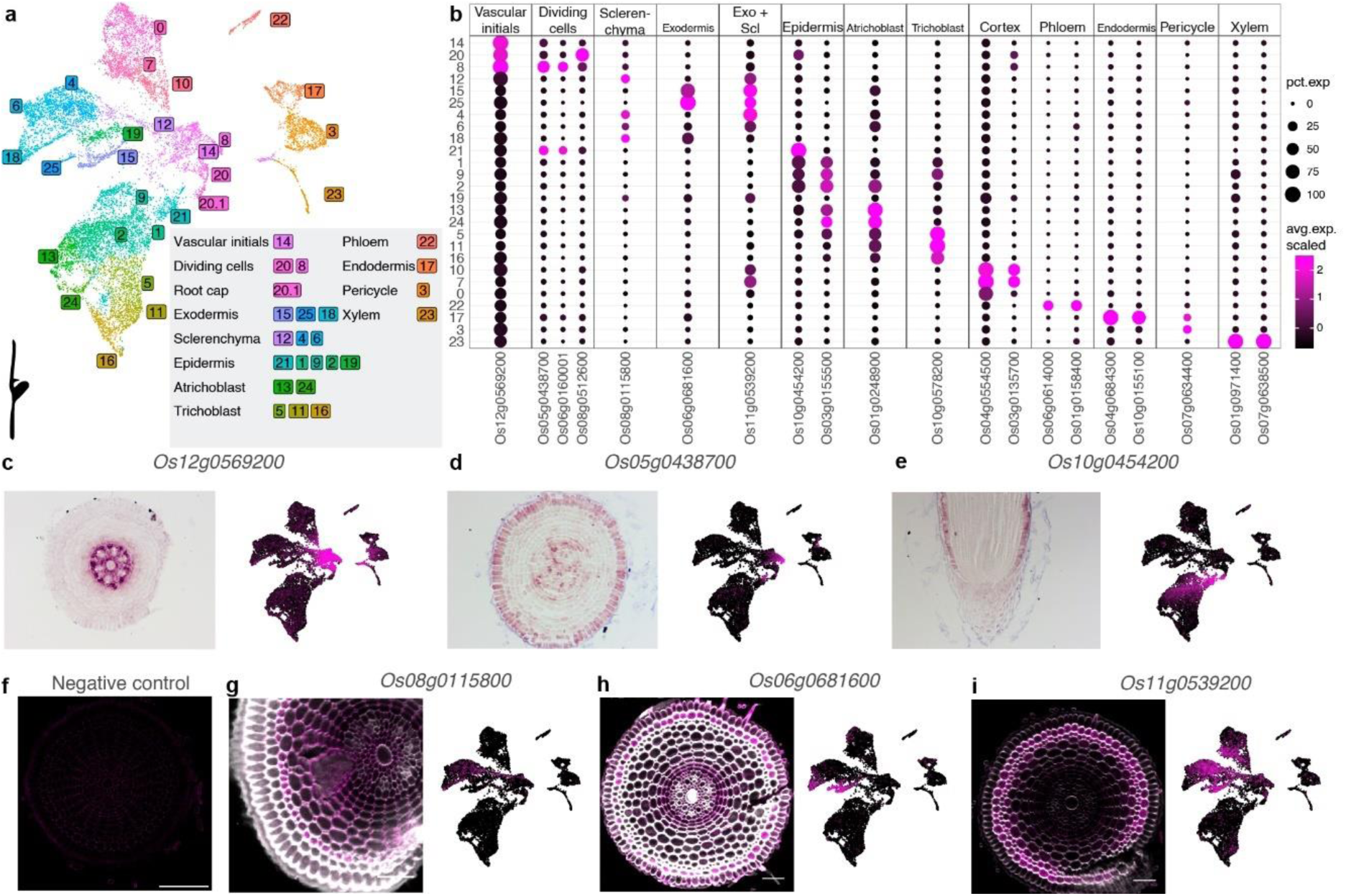
Cell type annotation of rice dataset. **a,** UMAP of merged mock samples from the rice dataset (R1, R2 and R3) showing the unsupervised clustering using the Louvain algorithm at a resolution of 1.2. Colors and labels represent distinct clusters and their matching cell type after manual annotation. **b,** Dot plot showing the average scaled expression of cell type specific marker genes previously described or validated in this study across the clusters of the rice reference dataset. **c-i,** RNA *in situ* hybridization showing the transcript localization of specific expressed genes within clusters in **a** that lacked clear annotation in cross sections of rice roots. **g-i,** Cell walls were stained with calcofluor white and represented in grey scale. **f,** Negative control without hybridization, showing unspecific binding of hairpin 488 to cell walls. Scale bars, 50 µM.

**Extended Data Fig. 6.**
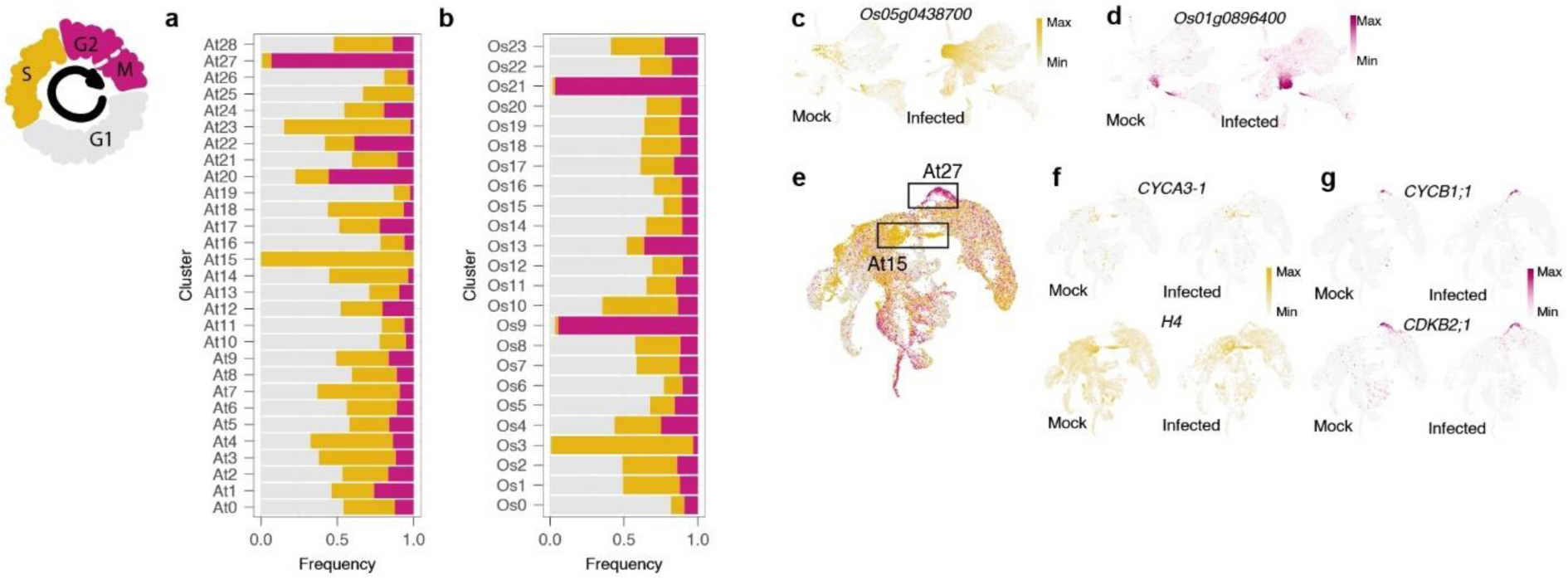
Cell phase annotation of *Arabidopsis* and rice datasets. **a,b,** Frequency of cells predicted in different cell cycle phases in *Arabidopsis* (**a**) and rice (**b**) datasets. **c,d,** UMAP showing the expression of marker genes for S-phase (**c**) and G2/M phase (**d**) in the rice dataset. **e,** UMAP highlighting the clusters enriched in cells predicted to be S-phase and G2/M-phase in the *Arabidopsis* dataset. **f,g,** UMAP showing the expression of marker genes for S-phase (**f**) and G2/M phase (**g**) in the *Arabidopsis* dataset.

**Extended Data Fig. 7.**
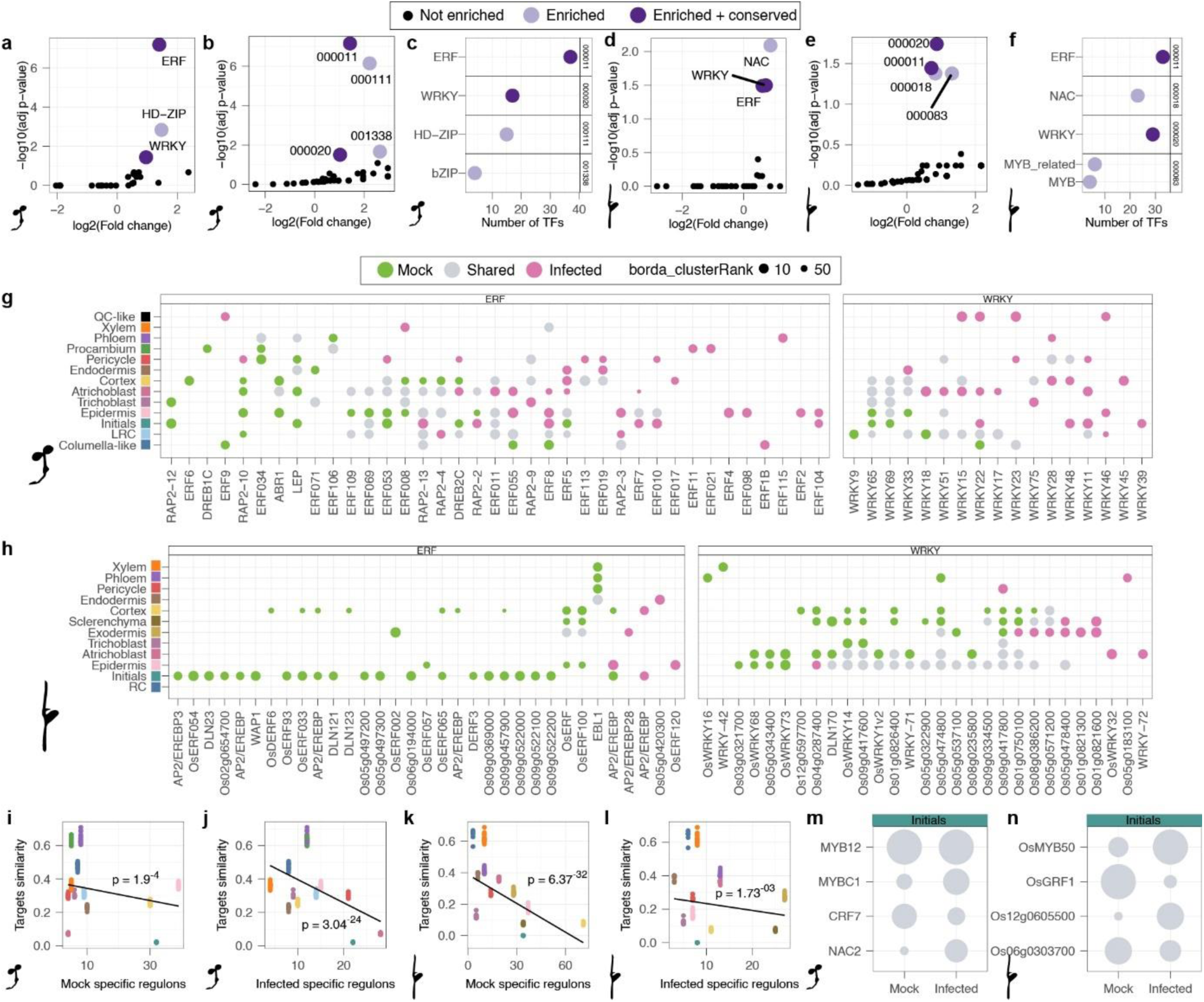
Analysis of TF families within the cell-type-specific regulons. **a-h,** Enrichment analysis of TF families and orthogroups within TFs that govern treatment-specific regulons in at least one cell type. **a,d,** Enrichment analysis of TF families in *Arabidopsis* (**a**) and rice (**d**) regulons. **b,e,** Enrichment analysis of orthogroups in *Arabidopsis* (**b**) and rice (**e**) regulons. **c,f,** Relation of enriched orthogroups with the TFs families that are contained in the orthogroups for *Arabidopsis* (**c**) and rice (**f**). **g,h,** Distribution of WRKY and ERF TFs and their corresponding regulons by cell type in the *Arabidopsis* (**g**) and rice (**h**) datasets. **i-n,** Analysis of TFs that govern regulons for both treatments within the same cell type. **i-l,** Correlation between the Jaccard index of targets similarity of TFs that are shared within cell types (grey bars in Fig. 5b,c) and the number of mock specific regulons (**i** and **k**) or infected specific regulons (**j** and **l**) in the same cell type, for *Arabidopsis* (**i** and **j**) and rice (**k** and **l**). The p-value of the correlation is shown in the plot. **m,n,** Best ranked shared regulons within initial cell types in *Arabidopsis* (**m**) and rice (**n**).

## Supplementary information

**Supplementary Table 1. Metrics, cluster analysis and evaluation of nematode responsive cells for Arabidopsis and rice datasets.**

**Supplementary Table 2. Gene orthology inference**

**Supplementary Table 3. Cell type and cell cycle phase annotation of Arabidopsis and rice datasets**

**Supplementary Table 4. Cell-type-specific gene regulatory networks of Arabidopsis and rice datasets**

**Supplementary Table 5. Information related to gene selection**

**Supplementary Table 6. Probes and oligonucleotide sequences related to in situ hybridization**

## References

1. Nicol, J. M. et al. Current Nematode Threats to World Agriculture. in Genomics and Molecular Genetics of Plant-Nematode Interactions (eds. Jones, J., Gheysen, G. & Fenoll, C.) 21–43 (Springer Netherlands, Dordrecht, 2011). doi:10.1007/978-94-007-0434-3_2.

2. Singh, S., Singh, B. & Singh, A. P. Nematodes: A Threat to Sustainability of Agriculture. Procedia Environmental Sciences 29, 215–216 (2015).

3. Wyss, U. & Grundler, F. M. W. Feeding behavior of sedentary plant parasitic nematodes. Netherlands Journal of Plant Pathology 98, 165–173 (1992).

4. Escobar, C., Barcala, M., Cabrera, J. & Fenoll, C. Chapter One - Overview of Root-Knot Nematodes and Giant Cells. in Advances in Botanical Research vol. 73 1–32 (2015).

5. Bird, A. F. THE ULTRASTRUCTURE AND HISTOCHEMISTRY OF A NEMATODE-INDUCED GIANT CELL. The Journal of Biophysical and Biochemical Cytology 11, 701–715 (1961).

6. Favery, B., Quentin, M., Jaubert-Possamai, S. & Abad, P. Gall-forming root-knot nematodes hijack key plant cellular functions to induce multinucleate and hypertrophied feeding cells. Journal of Insect Physiology 84, 60–69 (2016).

7. Kyndt, T., Vieira, P., Gheysen, G. & de Almeida-Engler, J. Nematode feeding sites: unique organs in plant roots. Planta 238, 807–818 (2013).

8. Jones, J. T. et al. Top 10 plant-parasitic nematodes in molecular plant pathology. Mol Plant Pathol 14, 946–961 (2013).

9. Bartlem, D. G., Jones, M. G. K. & Hammes, U. Z. Vascularization and nutrient delivery at root-knot nematode feeding sites in host roots. Journal of Experimental Botany 65, 1789–1798 (2014).

10. Caillaud, M.-C. et al. Root-knot nematodes manipulate plant cell functions during a compatible interaction. Journal of Plant Physiology 165, 104–113 (2008).

11. Goverse, A. & Mitchum, M. G. At the molecular plant-nematode interface: New players and emerging paradigms. Curr Opin Plant Biol 67, 102225 (2022).

12. Kayani, M. Z., Mukhtar, T. & Hussain, M. A. Effects of southern root knot nematode population densities and plant age on growth and yield parameters of cucumber. Crop Protection 92, 207–212 (2017).

13. Onkendi, E. M., Kariuki, G. M., Marais, M. & Moleleki, L. N. The threat of root-knot nematodes (eloidogyne spp.) in Africa: a review. Plant Pathology 63, 727–737 (2014).

14. Jain, R. K., Khan, Matiyar Rahaman & and Kumar, V. Rice root-knot nematode (Meloidogyne graminicola) infestation in rice. Archives of Phytopathology and Plant Protection 45, 635–645 (2012).

15. Yamaguchi, Y. L. et al. Root-Knot and Cyst Nematodes Activate Procambium-Associated Genes in Arabidopsis Roots. Front. Plant Sci. 8, (2017).

16. Olmo, R. et al. Root-knot nematodes induce gall formation by recruiting developmental pathways of post-embryonic organogenesis and regeneration to promote transient pluripotency. New Phytologist 227, 200–215 (2020).

17. Wyss, U. & Grundler, F. M. W. Feeding behavior of sedentary plant parasitic nematodes. Netherlands Journal of Plant Pathology 98, 165–173 (1992).

18. von Mende, N. Invasion and Migration Behaviour of Sedentary Nematodes. in Cellular and Molecular Aspects of Plant-Nematode Interactions (eds. Fenoll, C., Grundler, F. M. W. & Ohl, S. A.) 51–64 (Springer Netherlands, Dordrecht, 1997). doi:10.1007/978-94-011-5596-0_5.

19. Mitchum, M. G. et al. Nematode effector proteins: an emerging paradigm of parasitism. New Phytologist 199, 879–894 (2013).

20. Hewezi, T. & Baum, T. J. Manipulation of Plant Cells by Cyst and Root-Knot Nematode Effectors. MPMI 26, 9–16 (2013).

21. Zhao, J., et al. The root-knot nematode effector Mi2G02 hijacks a host plant trihelix transcription factor to promote nematode parasitism. Plant Comm 5, (2024).

22. Niu, J. et al. Msp40 effector of root-knot nematode manipulates plant immunity to facilitate parasitism. Sci Rep 6, 19443 (2016).

23. de Almeida Engler, J., et al. Molecular markers and cell cycle inhibitors show the importance of cell cycle progression in nematode-induced galls and syncytia. Plant Cell 11, 793–808 (1999).

24. Bird, A. F. THE ULTRASTRUCTURE AND HISTOCHEMISTRY OF A NEMATODE-INDUCED GIANT CELL. The Journal of Biophysical and Biochemical Cytology 11, 701–715 (1961).

25. Goverse, A. & Bird, D. The Role of Plant Hormones in Nematode Feeding Cell Formation. in Genomics and Molecular Genetics of Plant-Nematode Interactions (eds. Jones, J., Gheysen, G. & Fenoll, C.) 325–347 (Springer Netherlands, Dordrecht, 2011). doi:10.1007/978-94-007-0434-3_16.

26. Fosu-Nyarko, J., Jones, M. G. K. & Wang, Z. Functional characterization of transcripts expressed in early-stage Meloidogyne javanica-induced giant cells isolated by laser microdissection. Molecular Plant Pathology 10, 237–248 (2009).

27. Barcala, M. et al. Early transcriptomic events in microdissected Arabidopsis nematode-induced giant cells. The Plant Journal 61, 698–712 (2010).

28. Portillo, M. et al. Isolation of RNA from laser-capture-microdissected giant cells at early differentiation stages suitable for differential transcriptome analysis. Molecular Plant Pathology 10, 523–535 (2009).

29. Ramsay, K., Wang, Z. & Jones, M. G. K. Using laser capture microdissection to study gene expression in early stages of giant cells induced by root-knot nematodes. Molecular Plant Pathology 5, 587–592 (2004).

30. Ji, H. et al. Transcriptional analysis through RNA sequencing of giant cells induced by Meloidogyne graminicola in rice roots. J Exp Bot 64, 3885–3898 (2013).

31. Cabrera, J., Bustos, R., Favery, B., Fenoll, C. & Escobar, C. NEMATIC: a simple and versatile tool for the in silico analysis of plant–nematode interactions. Molecular Plant Pathology 15, 627–636 (2014).

32. Sijmons, P. C., Grundler, F. M. W., von Mende, N., Burrows, P. R. & Wyss, U. Arabidopsis thaliana as a new model host for plant-parasitic nematodes. The Plant Journal 1, 245–254 (1991).

33. Abril-Urias, P. et al. Divergent regulation of auxin responsive genes in root-knot and cyst nematodes feeding sites formed in Arabidopsis. Front. Plant Sci. 14, (2023).

34. Suzuki, R. et al. PUCHI Regulates Giant Cell Morphology During Root-Knot Nematode Infection in Arabidopsis thaliana. Front. Plant Sci. 12, (2021).

35. Bird, D. McK., Opperman, C. H. & Williamson, V. M. Plant Infection by Root-Knot Nematode. in Cell Biology of Plant Nematode Parasitism (eds. Berg, R. H. & Taylor, C. G.) 1–13 (Springer, Berlin, Heidelberg, 2009). doi:10.1007/978-3-540-85215-5_1.

36. Ozalvo, R. et al. Two closely related members of rabidopsis 13-lipoxygenases (13-LOXs), LOX3 and LOX4, reveal distinct functions in response to plant-parasitic nematode infection. Molecular Plant Pathology 15, 319–332 (2014).

37. Siddique, S., Wieczorek, K., Szakasits, D., Kreil, D. P. & Bohlmann, H. The promoter of a plant defensin gene directs specific expression in nematode-induced syncytia in Arabidopsis roots. Plant Physiology and Biochemistry 49, 1100–1107 (2011).

38. Lohar, D. P. et al. Cytokinins play opposite roles in lateral root formation, and nematode and Rhizobial symbioses. The Plant Journal 38, 203–214 (2004).

39. Mitchum, M. G. et al. The promoter of the Arabidopsis thaliana Cel1 endo-1,4-β glucanase gene is differentially expressed in plant feeding cells induced by root-knot and cyst nematodes. Molecular Plant Pathology 5, 175–181 (2004).

40. Zhou, W. et al. A Jasmonate Signaling Network Activates Root Stem Cells and Promotes Regeneration. Cell 177, 942–956.e14 (2019).

41. Grunewald, W. et al. A Role for AtWRKY23 in Feeding Site Establishment of Plant-Parasitic Nematodes. Plant Physiol 148, 358–368 (2008).

42. de Almeida Engler, J., et al. CCS52 and DEL1 genes are key components of the endocycle in nematode-induced feeding sites. The Plant Journal 72, 185–198 (2012).

43. Tosches, M. A. et al. Evolution of pallium, hippocampus, and cortical cell types revealed by single-cell transcriptomics in reptiles. Science 360, 881–888 (2018).

44. Guillotin, B. et al. A pan-grass transcriptome reveals patterns of cellular divergence in crops. Nature 617, 785–791 (2023).

45. Zhang, T.-Q., Chen, Y., Liu, Y., Lin, W.-H. & Wang, J.-W. Single-cell transcriptome atlas and chromatin accessibility landscape reveal differentiation trajectories in the rice root. Nat Commun 12, 2053 (2021).

46. Passalacqua, M. J. & Gillis, J. Coexpression enhances cross-species integration of single-cell RNA sequencing across diverse plant species. Nat. Plants 10, 1075–1080 (2024).

47. Barbie, D. A. et al. Systematic RNA interference reveals that oncogenic KRAS-driven cancers require TBK1. Nature 462, 108–112 (2009).

48. Ding, H., Blair, A., Yang, Y. & Stuart, J. M. Biological process activity transformation of single cell gene expression for cross-species alignment. Nat Commun 10, 4899 (2019).

49. Wendrich, J. R. et al. Vascular transcription factors guide plant epidermal responses to limiting phosphate conditions. Science 370, eaay4970 (2020).

50. Fernandez, A. et al. Transcriptional and Functional Classification of the GOLVEN/ROOT GROWTH FACTOR/CLE-Like Signaling Peptides Reveals Their Role in Lateral Root and Hair Formation. Plant Physiology 161, 954–970 (2013).

51. Haecker, A. et al. Expression dynamics of WOX genes mark cell fate decisions during early embryonic patterning in Arabidopsis thaliana. Development 131, 657–668 (2004).

52. Ferrari, C., Pérez, N. M. & Vandepoele, K. MINI-EX: Integrative inference of single-cell gene regulatory networks in plants. Molecular Plant 15, 1807–1824 (2022).

53. Staut, J., Pérez, N. M., Depuydt, T., Vandepoele, K. & Lukicheva, S. MINI-EX version 2: cell-type-specific gene regulatory network inference using an integrative single-cell transcriptomics approach. 2023.12.24.573246 Preprint at 10.1101/2023.12.24.573246 (2023).

54. Domínguez-Figueroa, J., Gómez-Rojas, A. & Escobar, C. Functional studies of plant transcription factors and their relevance in the plant root-knot nematode interaction. Front. Plant Sci. 15, (2024).

55. Warmerdam, S. et al. The TIR-NB-LRR pair DSC1 and WRKY19 contributes to basal immunity of Arabidopsis to the root-knot nematode Meloidogyne incognita. BMC Plant Biol 20, 73 (2020).

56. Chinnapandi, B. et al. Tomato SlWRKY3 acts as a positive regulator for resistance against the root-knot nematode Meloidogyne javanica by activating lipids and hormone-mediated defense-signaling pathways. Plant Signaling & Behavior 14, 1601951 (2019).

57. Chinnapandi, B., Bucki, Patricia & and Braun Miyara, S. SlWRKY45, nematode-responsive tomato WRKY gene, enhances susceptibility to the root knot nematode; M. javanica infection. Plant Signaling & Behavior 12, e1356530 (2017).

58. Kumar, A., Sichov, N., Bucki, P. & Miyara, S. B. SlWRKY16 and SlWRKY31 of tomato, negative regulators of plant defense, involved in susceptibility activation following root-knot nematode Meloidogyne javanica infection. Sci Rep 13, 14592 (2023).

59. Warmerdam, S. et al. Mediator of tolerance to abiotic stress ERF6 regulates susceptibility of Arabidopsis to Meloidogyne incognita. Molecular Plant Pathology 20, 137–152 (2019).

60. Dyer, S., et al. *Ethylene Response Factor* (*ERF*) genes modulate plant root exudate composition and the attraction of plant parasitic nematodes. International Journal for Parasitology 49, 999–1003 (2019).

61. Ribeiro, C. et al. The regeneration conferring transcription factor complex ERF115-PAT1 coordinates a wound-induced response in root-knot nematode induced galls. New Phytologist 241, 878–895 (2024).

62. Nahar, K., Kyndt, T., De Vleesschauwer, D., Höfte, M. & Gheysen, G. The Jasmonate Pathway Is a Key Player in Systemically Induced Defense against Root Knot Nematodes in Rice1[C]. Plant Physiol 157, 305–316 (2011).

63. Birnbaum, K. et al. A Gene Expression Map of the Arabidopsis Root. Science 302, 1956–1960 (2003).

64. Abril-Urias, P. et al. Divergent regulation of auxin responsive genes in root-knot and cyst nematodes feeding sites formed in Arabidopsis. Front. Plant Sci. 14, (2023).

65. Hutangura, P., Mathesius, U., Jones, M. G. K. & Rolfe, B. G. Auxin induction is a trigger for root gall formation caused by root-knot nematodes in white clover and is associated with the activation of the flavonoid pathway. Functional Plant Biol. 26, 221– 231 (1999).

66. Kyndt, T. et al. Redirection of auxin flow in Arabidopsis thaliana roots after infection by root-knot nematodes. Journal of Experimental Botany 67, 4559–4570 (2016).

67. Suzuki, R. et al. Local auxin synthesis mediated by YUCCA4 induced during root-knot nematode infection positively regulates gall growth and nematode development. Front. Plant Sci. 13, (2022).

68. Hu, Y. et al. OsHOX1 and OsHOX28 Redundantly Shape Rice Tiller Angle by Reducing HSFA2D Expression and Auxin Content. Plant Physiology 184, 1424–1437 (2020).

69. Possenti, M. et al. HD-Zip II transcription factors control distal stem cell fate in Arabidopsis roots by linking auxin signaling to the FEZ/SOMBRERO pathway. Development 151, dev202586 (2024).

70. Carabelli, M. et al. Arabidopsis HD-Zip II proteins regulate the exit from proliferation during leaf development in canopy shade. Journal of Experimental Botany 69, 5419– 5431 (2018).

71. Kyndt, T. Loss of susceptibility, an underexplored approach for durable resistance to plant-parasitic nematodes. Journal of Experimental Botany 74, 5422–5425 (2023).

72. Engler, J. de A., Favery, B., Engler, G. & Abad, P. Loss of susceptibility as an alternative for nematode resistance. Current Opinion in Biotechnology 16, 112–117 (2005).

73. Lee, L. R. et al. Glutathione accelerates the cell cycle and cellular reprogramming in plant regeneration. Developmental Cell 60, 1153–1167.e6 (2025).

74. Rusinque, L., Maleita, C., Abrantes, I., Palomares-Rius, J. E. & Inácio, M. L. Meloidogyne graminicola—A Threat to Rice Production: Review Update on Distribution, Biology, Identification, and Management. Biology 10, 1163 (2021).

75. Mantelin, S., Bellafiore, S. & Kyndt, T. Meloidogyne graminicola: a major threat to rice agriculture. Mol Plant Pathol 18, 3–15 (2016).

76. Wieczorek, K. et al. Expansins are involved in the formation of nematode-induced syncytia in roots of Arabidopsis thaliana. The Plant Journal 48, 98–112 (2006).

77. Guarneri, N. et al. Root architecture plasticity in response to endoparasitic cyst nematodes is mediated by damage signaling. New Phytologist 237, 807–822 (2023).

78. Olmo, R. et al. A Standardized Method to Assess Infection Rates of Root-Knot and Cyst Nematodes in Arabidopsis thaliana Mutants with Alterations in Root Development Related to Auxin and Cytokinin Signaling. in Auxins and Cytokinins in Plant Biology: Methods and Protocols (eds. Dandekar, T. & Naseem, M.) 73–81 (Springer, New York, NY, 2017). doi:10.1007/978-1-4939-6831-2_5.

79. Reversat, G., Boyer, J., Sannier, C. & Pando-Bahuon, A. Use of a mixture of sand and water-absorbent synthetic polymer as substrate for the xenic culturing of plant-parasitic nematodes in the laboratory. Nematol 1, 209–212 (1999).

80. De Rybel, B. et al. A Novel Aux/IAA28 Signaling Cascade Activates GATA23-Dependent Specification of Lateral Root Founder Cell Identity. Current Biology 20, 1697–1706 (2010).

81. Hirota, A., Kato, T., Fukaki, H., Aida, M. & Tasaka, M. The Auxin-Regulated AP2/EREBP Gene PUCHI Is Required for Morphogenesis in the Early Lateral Root Primordium of Arabidopsis. The Plant Cell 19, 2156–2168 (2007).

82. Xu, J. et al. A Molecular Framework for Plant Regeneration. Science 311, 385–388 (2006).

83. Tenorio Berrío, R., et al. Dual and spatially resolved drought responses in the Arabidopsis leaf mesophyll revealed by single-cell transcriptomics. New Phytologist 246, 840–858 (2025).

84. Turchi, L. et al. Arabidopsis HD-Zip II transcription factors control apical embryo development and meristem function. Development 140, 2118–2129 (2013).

85. Díaz-Manzano, F. E. et al. Long-Term In Vitro System for Maintenance and Amplification of Root-Knot Nematodes in Cucumis sativus Roots. Front. Plant Sci. 7, (2016).

86. Whitehead, A. G. & Hemming, J. R. A comparison of some quantitative methods of extracting small vermiform nematodes from soil. Annals of Applied Biology 55, 25–38 (1965).

87. McCarthy, D. J., Campbell, K. R., Lun, A. T. L. & Wills, Q. F. Scater: pre-processing, quality control, normalization and visualization of single-cell RNA-seq data in R. Bioinformatics 33, 1179–1186 (2017).

88. L. Lun, A. T., Bach, K. & Marioni, J. C. Pooling across cells to normalize single-cell RNA sequencing data with many zero counts. Genome Biology 17, 75 (2016).

89. Hao, Y. et al. Integrated analysis of multimodal single-cell data. Cell 184, 3573–3587.e29 (2021).

90. R Core Team. R: A language and environment for statistical computing. R Foundation for Statistical Computing (2021).

91. Jammes, F. et al. Genome-wide expression profiling of the host response to root-knot nematode infection in Arabidopsis. The Plant Journal 44, 447–458 (2005).

92. Meijer, A. et al. Dicer-like 3a mediates intergenerational resistance against root-knot nematodes in rice via hormone responses. Plant Physiology 193, 2071–2085 (2023).

93. Atighi, M. R., Verstraeten, B., De Meyer, T. & Kyndt, T. Genome-wide shifts in histone modifications at early stage of rice infection with Meloidogyne graminicola. Molecular Plant Pathology 22, 440–455 (2021).

94. Desmedt, W. et al. Rice diterpenoid phytoalexins are involved in defence against parasitic nematodes and shape rhizosphere nematode communities. New Phytologist 235, 1231–1245 (2022).

95. Van Bel, M. et al. PLAZA 5.0: extending the scope and power of comparative and functional genomics in plants. Nucleic Acids Res 50, D1468–D1474 (2021).

96. Buchfink, B., Reuter, K. & Drost, H.-G. Sensitive protein alignments at tree-of-life scale using DIAMOND. Nat Methods 18, 366–368 (2021).

97. Enright, A. J., Van Dongen, S. & Ouzounis, C. A. An efficient algorithm for large-scale detection of protein families. Nucleic Acids Res 30, 1575–1584 (2002).

98. Emms, D. M. & Kelly, S. OrthoFinder: phylogenetic orthology inference for comparative genomics. Genome Biology 20, 238 (2019).

99. Price, M. N., Dehal, P. S. & Arkin, A. P. FastTree 2 – Approximately Maximum-Likelihood Trees for Large Alignments. PLoS One 5, e9490 (2010).

100. Katoh, K. & Standley, D. M. MAFFT Multiple Sequence Alignment Software Version 7: Improvements in Performance and Usability. Mol Biol Evol 30, 772–780 (2013).

101. Chen, K., Durand, D. & Farach-Colton, M. NOTUNG: a program for dating gene duplications and optimizing gene family trees. J Comput Biol 7, 429–447 (2000).

102. Di Tommaso, P. et al. Nextflow enables reproducible computational workflows. Nat Biotechnol 35, 316–319 (2017).

103. Mercatelli, D., Lopez-Garcia, G. & Giorgi, F. M. corto: a lightweight R package for gene network inference and master regulator analysis. Bioinformatics 36, 3916–3917 (2020).

104. Zöllner, N. R. et al. An RNA *in situ* hybridization protocol optimized for monocot tissue. STAR Protocols 2, 100398 (2021).

105. Bolger, A. M., Lohse, M. & Usadel, B. Trimmomatic: a flexible trimmer for Illumina sequence data. Bioinformatics 30, 2114–2120 (2014).

106. Langmead, B. & Salzberg, S. L. Fast gapped-read alignment with Bowtie 2. Nat Methods 9, 357–359 (2012).

107. Lawrence, M. et al. Software for Computing and Annotating Genomic Ranges. PLOS Computational Biology 9, e1003118 (2013).

108. Love, M. I., Huber, W. & Anders, S. Moderated estimation of fold change and dispersion for RNA-seq data with DESeq2. Genome Biology 15, 550 (2014).

